# Non-Equilibrium Snapshots of Ligand Efficacy at the μ-Opioid Receptor

**DOI:** 10.1101/2025.05.26.656223

**Authors:** Michael J. Robertson, Makaía M. Papasergi-Scott, Maria Claudia Peroto, Balazs R. Varga, Susruta Majumdar, Georgios Skiniotis

## Abstract

Distinct ligands for the same G-protein coupled receptor (GPCR) activate intracellular signaling partners to varying extents, but the molecular mechanisms driving these differences remains elusive. Hypothesizing that such differences in signaling efficacy may be captured structurally in intermediate states under non-equilibrium conditions, we implemented a time-resolved (TR) cryo-EM approach to visualize the GTP-induced activation of the Gαiβγ heterotrimer by the μ-opioid receptor (MOR) bound to three ligands displaying partial, full, or super-agonism on the receptor. We resolved ensembles of conformational states along the G-protein activation pathway, including a previously unobserved intermediate state that enabled us to visualize receptor dynamics as a function of bound ligand. The results demonstrate ligand-dependent differences in state occupancy and conformational stability, with higher ligand efficacy correlating with increased dynamics of the receptor’s transmembrane (TM) helices 5 and 6. Furthermore, we identify key mechanistic differences in the GTP-induced activation of Gi compared to Gs that likely underlie their distinct activation kinetics. Corroborated by molecular dynamics (MD) simulations, these findings provide a dynamic structural landscape of GPCR-G-protein interactions for ligands of different efficacy and suggest partial agonists may produce a ‘kinetic trap’ during G-protein activation.

## Introduction

GPRCs represent an outstanding class of therapeutic targets^1^, with a strong need for next-generation medicines with expanded therapeutic windows and reduced adverse effects. One of the most pursued mechanisms for achieving this goal is the development of agonists that precisely tune the receptor’s activity towards signal transducers such as G-proteins. In parallel, there are efforts to mitigate negative side effects by identifying GPCR ligands that drive limited activation of specific signaling pathways while maintaining or increasing their efficacy at others, an effect referred to as “biased signaling” or “signaling specificity”^2^.

An obstacle in designing bespoke GPCR ligands with tunable activity is the lack of mechanistic understanding of how differing degrees of agonism are transmitted intracellularly to transducers. In pharmacological assays, amplitudes of downstream signaling by an agonist are commonly compared to that of an endogenous or reference agonist, with those producing a comparable degree of signaling classified as ‘full’ agonists, those achieving less activation considered ‘partial’ agonists, and those achieving significantly greater activation designated as ‘super’ agonists.

Despite case studies of numerous ligands with various pharmacological profiles bound to the same receptor-G-protein complex^3-6^, the mechanisms by which a partial, full, or a super agonist transduce unique “signals” to tune G-protein activation remain unclear. Indeed, available structures of GPCR-G-protein complexes reveal detailed differences in ligand recognition but show virtually identical conformations for the intracellular side of the receptor, regardless of the bound ligand identity, while the nucleotide-depleted G-protein is also observed in the same state regardless of receptor.

A key question concerning the concept of efficacy relates to the receptor conformations promoted by partial versus full agonists. One possibility is that partial and full agonists promote different receptor conformational states. In this model, receptor-G-protein complexes progress from initial engagement to nucleotide release through two or more alternate pathways. These pathways would have different intermediate conformational waypoints, and pathways promoted by full agonists would be more efficient than the pathways promoted by partial agonists. An alternative possibility is that partial and full agonists promote the same conformational changes but do so at different rates. In this model, receptor-G-protein complexes progress through a single pathway with a predetermined set of intermediate conformational waypoints, but full agonists would be better than partial agonists at lowering energy barriers between waypoints. The two models are not mutually exclusive, but distinguishing their contributions in agonism has not been possible with traditional structural biology approaches. Crystal structures of nanobody-bound GPCRs^7,8^ and spectroscopic studies^9^ have shed some light on how ligands impact intracellular receptor conformations, showing that distinct, ligand-dependent receptor conformations do exist and drive pharmacology. However, capturing such conformations at high resolution has been challenging due to the inherently dynamic process of G-protein activation by a GPCR, which involves G-protein association steps followed by exchange of GDP to GTP and subsequent dissociation from the receptor.

Here we use a time-resolved (TR) cryo-EM approach to compare three agonists, with widely differing efficacies, bound to a receptor for which agonist efficacy has vital clinical relevance. The μ-opioid receptor (MOR) is the target of morphine and other opioids that remain highly effective analgesics^11^ but have fueled an ongoing health crisis due to their addictive potential combined with tolerance and respiratory depression^12^. Evidence suggests there may be room to optimize the safety window of opioids by modulating their signaling properties^13-15^, but a significantly improved, FDA-approved opioid currently remains elusive. We assess the GTP-induced activation of inhibitory Gi bound to the MOR activated by three agonists: DAMGO (D-Ala^2^, N-MePhe^4^, Gly-ol-enkephalin), a synthetic analogue of enkephalin that represents a canonical agonist; mitragynine pseudoindoxyl (MP), an active metabolite of kratom plant mitragynine that shows partial agonism; and lofentanil (LFT), a synthetic fentanyl-based ligand with such high potency, efficacy, and duration of action that is regarded as a MOR super-agonist too dangerous for even veterinary use in large mammals.

We resolve several distinct intermediate ensembles of GTP-triggered MOR-Gi states displaying various extents of G-protein activation. These include ordering of the linker region between the RHD and AHD upon GTP binding; semi-closure of the AHD and initial rearrangement of α5 and its receptor interface; full closure of the AHD and total withdrawal of α5 from the receptor; and finally, rearrangement of switch II and switch III loops driving several angstrom separation between Gα and Gβγ. Notably, we observe ligand-dependent differences in both the relative populations of particles in each ensemble and the conformational dynamics underlying each structural state. Intermediate states are sampled differently in a manner consistent with full agonists promoting more efficient progress along a common activation pathway. Supported by molecular dynamics (MD) simulations, our findings point to full agonist-bound receptors being more dynamic, allowing rapid progress past low energy conformational traps that slow G-protein activation by partial agonists.

## Results

### Structural ensembles of Nucleotide-Free and GTP-bound MOR-Gi

Previous cryo-EM work with MOR-Gi leveraged a combination of modified G-protein and/or immunoglobulin fragments to stabilize the assembly, thereby potentially altering the dynamics of the nucleotide-free complex. To our knowledge, the only exception is our prior structure of MP-bound MOR-Gi that employed a wildtype Gi heterotrimer without antibody fragments^6^. In a first step, to assess the conformational distribution of the AHD in the MP bound MOR-Gi complex, we applied 3D variability analysis (3DVA)^16^, an approach that identifies the principal directions of conformational differences in a cryo-EM dataset and generates successive reconstructions along these coordinates. The 3DVA analysis of MP bound MOR-Gi indicates that the alpha helical domain (AHD) occupies a rather limited range of open states while loosely bound on Gβγ without any closed-like states observed (Extended Data Figure 1A). This contrasts the behavior of the stimulatory Gαs, probed in our earlier cryo-EM study with the β_2_-adrenergic receptor, which showed that the AHD has intrinsically high dynamics, opening up and closing against the Ras homology domain (RHD) even in absence of nucleotide.

To probe if there is a ligand dependence to AHD positioning, we obtained additional cryo-EM structures of wildtype MOR-Gi heterotrimer with DAMGO (2.8 Å) and LFT (2.5 Å) (Figure 1b, Extended Data Figure 1a,2a). The ligand is well-resolved in each case (Figure 1c), and 3DVA for each condition reveals largely the same AHD positioning as in the MP data (Extended Data Figure 3a). Thus, within what can be resolved for this dynamic component under our experimental conditions, the behavior of the somewhat flexible but overall localized open-state AHD in a nucleotide-free Gi protein appears to be independent of whether a partial, full, or super agonist is bound to the receptor. This preference for an open AHD in the nucleotide depleted state is supported by 1 μs MD simulations of the MOR-Gi complex. Starting from a closed AHD state, 2 out of 5 trajectories transition to the open state and remain open, while all 5 simulations started from the AHD-open state remain open throughout the simulation run (Extended Data Figure 3b). These results suggest that, in contrast to Gs^10^, the AH-closed state for Gi is markedly higher in energy than the open states in the absence of nucleotide, and that the barrier between the two is likely significant. This is consistent with both our cryo-EM results and prior DEER studies of the rhodopsin-Gi complex^17^, which demonstrated that the AHD of Gi in the active rhodopsin-Gi complex is virtually all in an open state in the absence of nucleotide.

**Figure 1:**
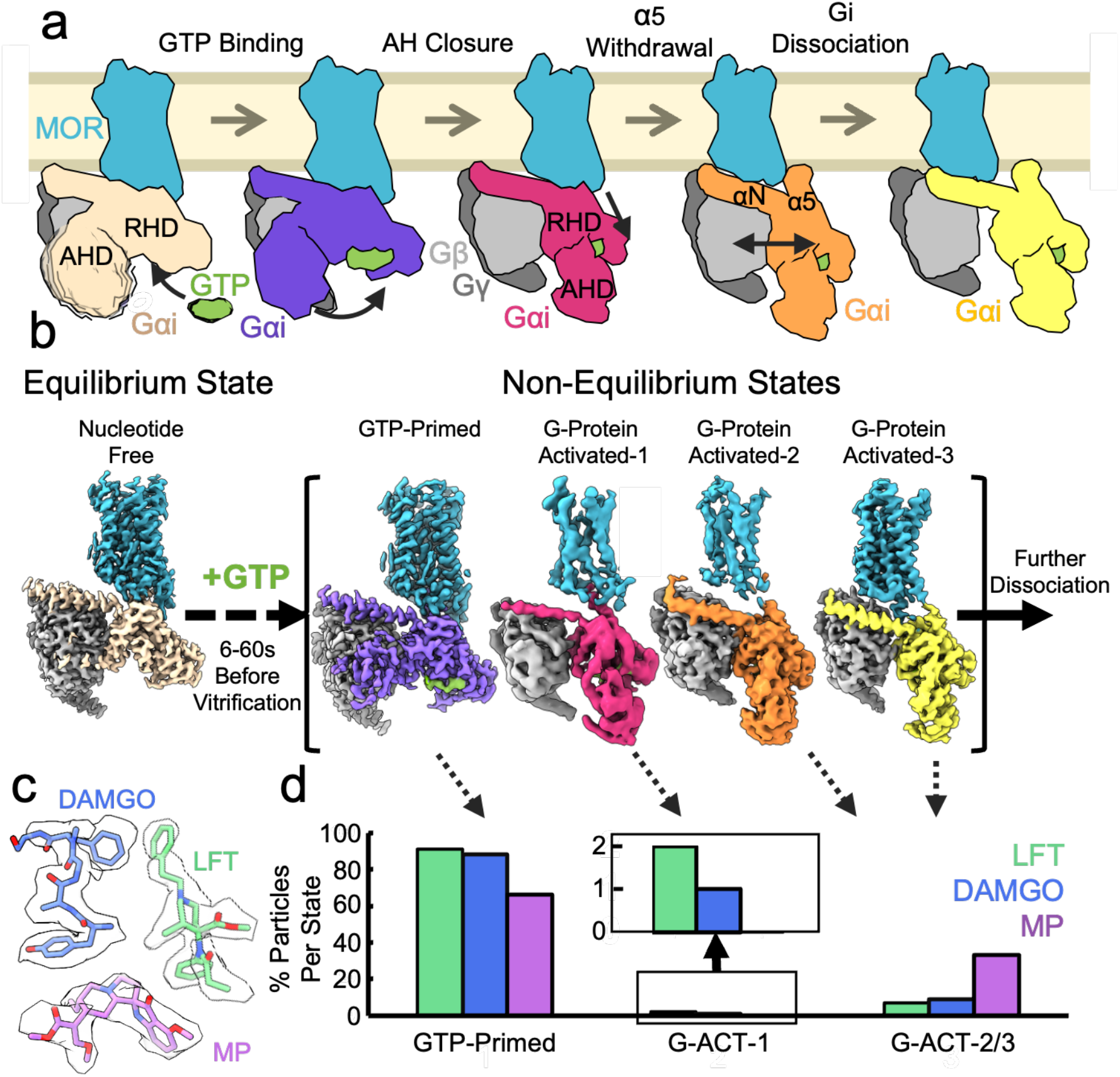
Conformational Dynamics of Gi Activation. **a)** Schematic of the process of GTP-induced activation of G-protein heterotrimer from MOR. Binding of the GTP induces AHD closing, which results in a withdrawal of the α5 helix from the receptor and subsequent dissociation of the G-protein subunits from each other **b)** Maps of the major states resolved in this work. After addition of GTP, four primary states are observed: an AHD opened but ordered state, termed ‘GTP-Primed’; a loosely AHD closed state, termed ‘G-ACT-1’, where the α5 helix remains partly engaged in the receptor; a tightly AHD closed state, termed ‘G-ACT-2’, where the G-protein is fully disengaged from the canonical intracellular pocket of the receptor; and a state termed ‘G-ACT-3’ where the G-proteins have begun to separate. **c)** The three ligands used in this work, the canonical full agonist DAMGO, the partial agonist MP, and the super agonist LFT shown in their cryo-EM density map from the GTP-Primed state. **d)** A plot of the relative particle populations in each of the major GTP-bound ensembles resolved with each of the three ligands in this work, with an inset of the percent population of particles in the G-ACT-1 state.

To visualize the GTP-induced activation of Gi1 from the MOR, we implemented TR cryo-EM to examine pre-steady state intermediate complexes between Gi and MOR bound to the three ligands vitrified 6s to 60s after adding GTP to the preformed nucleotide-free complex. The experimental concept and execution is similar to our recent work with Gs activation by the β_2_AR^10^, but with incubation with GTP leading to dissociation of the Gi protein from MOR. Here we were able to resolve three overarching ensembles of states, as observed during early stages of 3D classification (Extended Data Figure 1b).

The predominant state of the complex, regardless of timepoint or ligand, is an AHD-open state with well-resolved density for bound GTP (Figure 1b,d; Extended Data Figure 2a), referred to here as a ‘GTP-Primed’ state. However, unlike the nucleotide-free open state, the linker between the Ras Homology Domain (RHD) and AHD, as well as the AHD itself, undergo a substantial ordering against Gβγ (Extended Data Figure 3a,c). Minor filtering of particles based on the 3DVA analysis to select those with the most stable positioning facilitated reconstructions to 2.8-3.0 Å globally, with high quality density for the AHDs in the open state. (Extended Data Figure 2b). These structures reveal that the Gα AHD/Gβ interface involves relatively sparse, primarily hydrophilic interactions, including hydrogen bonding between Y61 and Y69 of Gα and R96 of Gβ (Extended Data Figure 3d), consistent with the lack of tight binding of the AHD to Gβγ when the linker regions are not ordered by nucleotide.

The second conformational state identified shows a closed-like AHD that is not yet fully ordered against the RHD. In this state the α5 helix of the G-protein is largely still extended into the intracellular cavity of an active-like receptor, as TM5 and TM6 are clearly swung outward (Figure 1b, Extended Data Figure 2a), a state we refer to as G-protein activation intermediate conformation 1 (G-ACT-1). Although G-ACT-1 is the lowest resolution state (due to the very small population of particles, regardless of ligand), the overall map for G-ACT-1 is of sufficient quality to dock G-protein and receptor models. Local refinement of G-ACT-1 on the G-protein produces map features that facilitated modeling of the protein backbone and many large side chains. This state shows interactions highly suggestive of an early intermediate state in G-protein activation, with some space remaining between the AHD and RHD, as well as partial ordering of many of the key switches I-III of the G-protein around GTP.

The third conformational ensemble identified, encompassing states we refer to as G-protein activation intermediate conformations 2 and 3 (G-ACT-2/3), differs from G-ACT-1 in that the AHD has fully closed against the RHD and the α5 helix of the G-protein has completely withdrawn from the receptor (Figure 1b). 3DVA-based particle sorting on this state with MP and DAMGO enabled us to identify two distinct G-protein states: G-ACT-2, with the typical arrangement of Gα and Gβγ in cryo-EM structures^6,10^, and G-ACT-3, where Gα has partially separated from Gβγ, which, to our knowledge, has not been observed before. Notably, the G-ACT-2/3 states reveal ligand-dependent differences in receptor dynamics (Figure 2a, Extended Data Figure 2a): In MP bound MOR, we observe a well ordered and readily modellable receptor density; in DAMGO-bound MOR the results show a partially dynamic receptor density; and with LFT we observe a highly disordered receptor density. On the other hand, for all three ligands, local refinement of the G-protein produces a map with sufficient quality to allow for confident backbone modeling (Extended Data Figure 2b).

**Figure 2:**
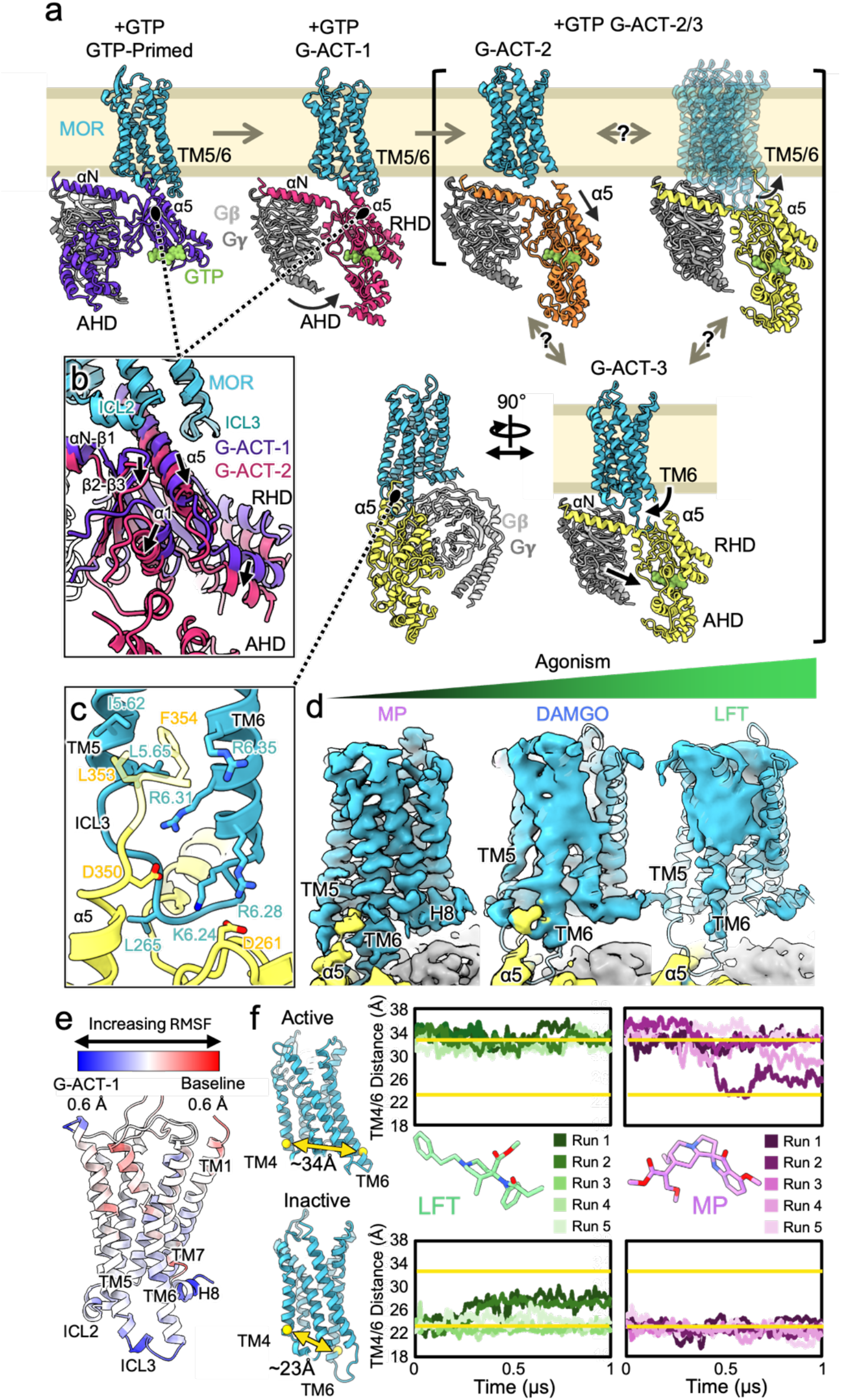
Ligand-Dependent Differences in MOR-Gi Complex Activation. **a)** Structures of the states resolved in this work. A representation of the ensemble of states observed in the G-ACT-2/3 class with all three ligands is shown in brackets. This includes G-ACT-2 where Gα and βγ are together, G-ACT-3 where Gα and βγ have begun to separate, and a third ‘ensemble’, not strictly mutually exclusive from G-ACT-2/3, where the poorly resolved receptor is largely dynamic and unengaged from the G-protein. Which states are stabilized and how well resolved the receptor is depends on the ligand (Extended Data Figure 2a). **b)** Inset comparison of the G-ACT-1 and G-ACT-2 states aligned on the MOR showing the ∼1-3 Å drop of Gαi relative to the receptor. **c)** Inset showing the interface of the MOR TM5-ICL3-TM6 region (teal) and Gαi (yellow) in the G-ACT-3 state with MP. The semi-transparent region reflects areas where the cryo-EM map has features but of insufficient resolution for confident modeling. **d)** Comparison of the MOR-Gi interface in the G-ACT-3 state, where the 3DVA particle sorting also reflects the most stable receptor conformation. Maps are presented for MP, DAMGO, and LFT highlighting the loss in resolution of the base of TM5 and TM6 with increasing potency of the agonist. **e)** Average difference in RMSF between baseline and G-ACT-1 simulations. **f)** MD plots showing TM4-TM6 distance starting from active (top) and inactive (bottom) receptors with LFT and MP. Yellow lines indicate active vs inactive TM4-6 distance, with the measured distances denoted on cartoons of MOR in inactive and active states.

The overall populations of the three ensembles of MOR-Gi states vary significantly with the three ligands (Figure 1d): LFT shows the highest percentage of the population in the GTP-Primed (∼90%), with very little G-ACT-1 (∼2%) and modest G-ACT-2/3 (∼8%) complexes. MP, by contrast, has almost no G-ACT-1, but a substantial enrichment of G-ACT-2/3 (∼33%) of complexes. Notably, the relative populations of these states with DAMGO falls in between those of the super-agonist LFT and the partial agonist MP, revealing a consistent trend of decreasing GTP-Primed and G-ACT-1 complexes with increasing G-ACT-2/3 complexes as agonist efficacy becomes progressively lower.

In comparing the relative populations of the three major ensembles for LFT and DAMGO across different timepoints, we could not identify a statistically significant trend. The low occupancies of the G-ACT-1 states compounded with limitations in classification result in error bars on particle assignments that, while allowing for comparing these populations across different ligands, preclude us from monitoring the progression of states at sequential time points within the time-scale of our experiment (6-60 sec). Nevertheless, the three conformational ensembles appear to provide sequential snapshots of G-protein activation by MOR, as also compared to our previous work with β_2_AR-Gs^10^. We can order these states along the activation pathway, as judged by the relative positioning of the AHD domain, the state of the C-terminal helix α5, and the rearrangements of several key microswitches within the G-protein, discussed in greater detail in subsequent sections (Supplementary Video 1). We also observe 2D/3D classes in the presence of GTP that appear to represent MOR-Gβγ complexes with largely unresolvable Gα density, which we speculate may represent a state even further along the G-protein activation pathway (Extended Data Figure 1b).

### Ligand-Dependent Receptor Dynamics During G-protein Activation

Models can be confidently built or docked for the majority of these conformational states for each of the ligands (Figure 2a), allowing for a ligand-dependent comparison of the receptor behavior throughout the G-protein activation cycle. As the complex transitions from the GTP-Primed state to the G-ACT-1 state and the AHD begins to close, the receptor remains in an active state-like conformation, with an extended TM6, and the G-protein is still engaged with the intracellular core of the receptor (Extended Data Figure 2a). However, we observe a roughly 1-3 Å downward shift of the RHD away from MOR, revealing a substantial weakening of receptor-G-protein interactions (Figure 2b). Examination of map densities suggests that the α5 helix has started to retract towards the G-protein, with parallel disruption of interactions between the αN-β1 loop and intracellular loop 2 (ICL2) of the receptor, and repositioning of ICL3 away from the α5 helix as it disrupts its interactions with the G-protein. These observations suggest that GTP binding drives rearrangements of both the C-terminal α5 helix, through the nucleotide’s guanine base interactions, but also the αN-β1 loop, through the nucleotide’s phosphate interactions with the β1-β2 loop. The weakening of these two central interactions initiates the uncoupling between receptor and G-protein further destabilizing ICL3 interactions. In support of this mechanism, MD simulations comparing the internal root mean squared fluctuations (RMSF) in the receptor with the nucleotide-free baseline and G-ACT-1 LFT-bound state reveal increased dynamics in the G-ACT-1 state in ICL3 and helix 8, and to a lesser extent ICL2 (Figure 2e).

In contrast to G-ACT-1, the α5 helix of the G-protein in the G-ACT-2/3 states has shifted entirely out of the MOR intracellular cavity and has fully retracted downward towards the nucleotide, allowing the receptor to deviate from its position over the G-protein. In the G-ACT-2/3 state with MP, we observe that the majority of the particles have a well-ordered receptor with the TM6 helix shifted inward, representing an inactive state^8^, wedging itself between Gα and Gβγ subunits. 3DVA analysis shows that the main source of receptor variability in this state is due to a minor propensity of the receptor to separate from the G-protein (Extended Data Figure 2a). A reconstruction from the most stable ‘inactive-like’ receptor states (corresponding to G-ACT-3) yields a 3.6 Å map where the receptor can be modeled reliably, including much of ICL3 (Figure 2d, Extended Data Figure 2a,4a). In this state, TM5, ICL3, and TM6 fall into the cleft between the C-terminus/RHD of Gα and the back of the β-barrel of Gβγ. While the resolution for the very tip of the C-terminus of Gα is poor (Extended Data Figure 4a), modeling suggests that the sidechain of F354 is hydrophobically packed against residues I5.62 and L5.56 (Ballesteros-Weinstein numbering system^18^) between the outer sides of TM5 and TM6. Several cationic residues including R6.35, R6.31, R6.25, and K6.24 are in proximity to interact with the anionic C-terminal carboxylate, D350 of the α5 helix, and also D261 of Gα (Figure 2c). Notably, the interaction of Gα with MOR at this interface mimics that of megabody 6 (Mb6) obtained with the inactive-state receptor in complex with an antagonist (PDB:7UL4, Extended Data 4b)^19^, particularly W489 and several anionic residues of the megabody. A survey of family A GPCRs reveals that this polycationic outward face at the base of TM5-ICL3-TM6 is found broadly across most receptors (Extended Data Figure 4c,d) and thus this may represent a general mechanism by which transient interactions can form between Gi and family A GPCRs. Several of the G-protein residues are also conserved, with D350 either Asp or Glu in most Gα and D261 also anionic in G12/13, suggesting this intermediate may exist with other Gα. While our structure derives from our attempts to image the GTP-induced dissociation of Gi from MOR, this state shares interactions with a pre-coupled/intermediate state identified by recent MD simulations examining β_2_AR-Gs association, which highlighted a similar role for these outward-facing cationic receptor residues (20).

3DVA of the MP-bound complexes in the G-ACT-2/3 state revealed conformational heterogeneity within the G-protein heterotrimer, with 3D reconstructions showing a ∼4-7 Å displacement of Gα away from Gβγ (Figure 2a, 3g, Extended Data 2a, Supplementary Video 2). Of note, the extent of separation between the Gi subunits correlated with the stability of receptor densities. Similar heterogeneity in terms of both receptor and G-protein were observed for DAMGO (Extended Data Figure 2a, Supplementary Video 3), but with worse resolution for the receptor region. The loss in resolvability for DAMGO-bound receptor densities in the G-ACT-2/3 state is particularly prominent around the cytoplasmic side of TM5 and to a lesser extent TM6, even when we select for the particles with the most stable receptor (Figure 2d). This likely reflects increased receptor dynamics and not lower data quality, as local refinement of the G-protein for DAMGO G-ACT-2/3 still reaches 3.6 Å global resolution (Extended Data Figure 2b). The trend of increased receptor dynamics is even more prominent with LFT, where a single receptor state cannot be resolved, especially at the cytoplasmic ends of TM5 and TM6 (Figure 2d, Supplementary Video 4). A recent DEER study has suggested that MOR in the presence of MP occupies a primarily inward state of TM6, while DAMGO produces increasing populations of open-state TM6 and LFT-bound MOR occupies almost entirely this fully TM6 open conformation^9^. This is consistent with our results, where the inward TM6 conformation found in G-ACT-2/3 is most prominently observed with MP. By contrast, little, if any, LFT-bound receptors are in this G-protein engaged but inactive-like conformation, whereas DAMGO appears to drive an intermediate state between MP and LFT, with both inward and outward configurations of TM6. We further probed this hypothesis with MD simulations of the MOR alone, starting from both the inactive and active states bound to LFT or MP (Figure 2f). In just a single microsecond, 2 out of 5 trajectories of LFT-bound MOR starting from the inactive state had begun to transition to an active-like state with opening of TM6 (as measured by TM4-TM6 distance analogous to Jiawe *et al*.^9^), while all the active-like trajectories of LFT remained open (Figure 2f). In contrast, in 2 out of 5 of the active-like trajectories of MP we observe TM6 return towards a closed conformation, while all the inactive-like trajectories remain with TM6 in an inward conformation (Figure 2f). Together these results further support the structural differences and analysis of conformations underlying partial, full, and super-agonism at MOR.

**Figure 3:**
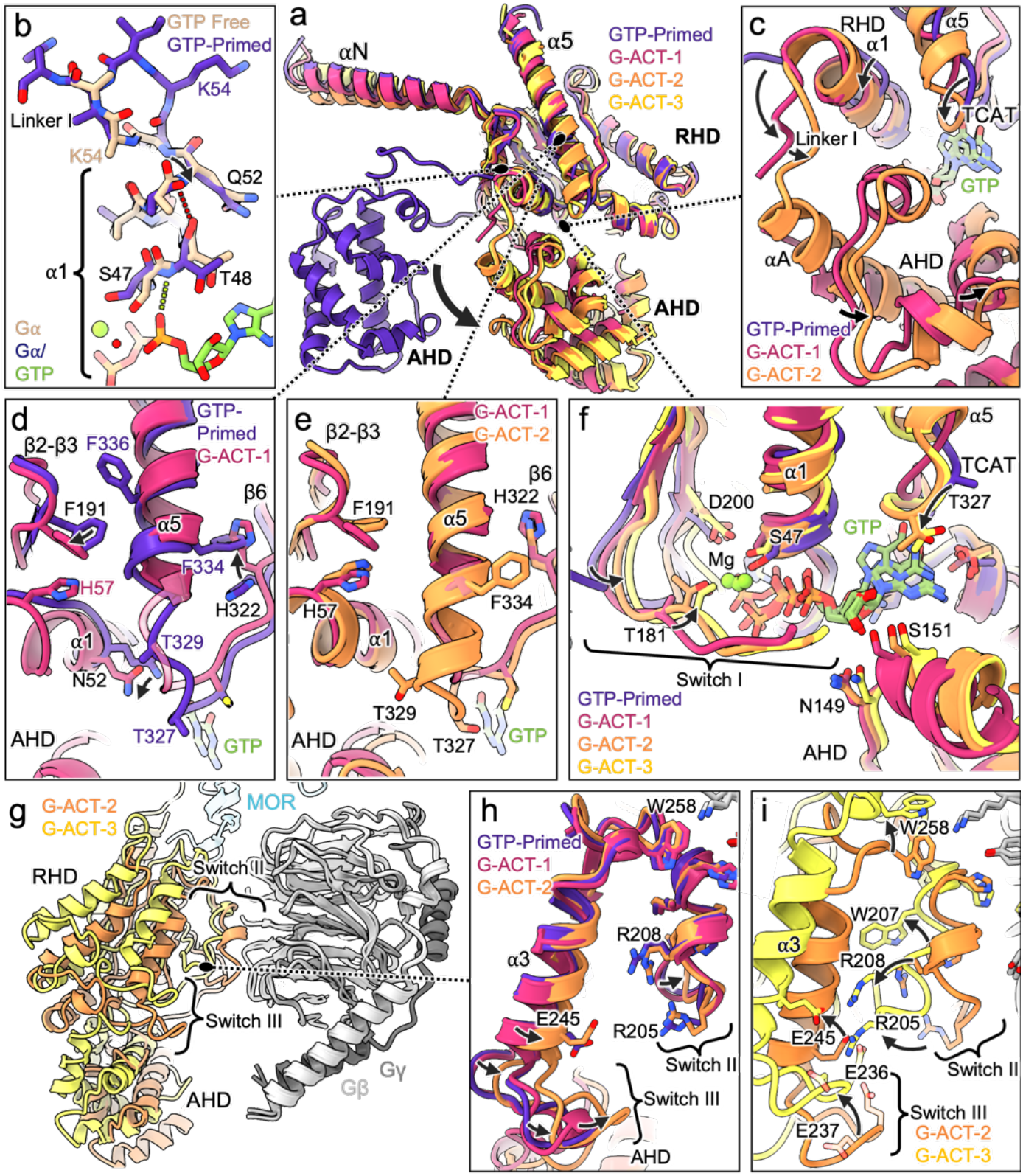
GTP-Induced Rearrangement of Gi Microswitches. **a)** Aligned structures of Gi heterotrimer in GTP-Primed (purple), LFT G-ACT-1 (magenta), MP G-ACT-2 (orange), and MP G-ACT-3 (yellow) states. **b)** Inset of the GTP-Primed and nucleotide-free state (tan) highlighting the structural shifts in nucleotide binding leading to Linker I ordering. **c)** Inset of the Linker I and AHD closing and ordering between the GTP-Primed, LFT G-ACT-1, and MP G-ACT-2 states. **d)** Inset showing the changes in the α1 helix, β2-β3 loop, and β6 with respect to the α5 helix in transitioning from the GTP-Primed state to the LFT G-ACT-1 state. **e)** Inset showing the changes in the α1 helix, β2-β3 loop, and β6 with respect to the α5 helix in transitioning from the LFT G-ACT-1 state to MP G-ACT-2 state. **f)** Inset highlighting the gradual closing and ordering of Switch I-Linker II when transitioning between all four states of the G-protein heterotrimer. **g)** MP G-ACT-2 and MP G-ACT-3 aligned on Gβγ. **h)** Inset of the changes in the Switch II, Switch III, and α3 regions of the GTP-Primed, LFT G-ACT-1, and MP G-ACT-2 states. **i)** Inset of the changes in the Switch II, Switch III, and α3 regions of the GTP-bound MP G-ACT-2 and MP G-ACT-3 states.

### Gi1 Transitions upon GTP-Induced Activation

These structures allow us to track crucial intermediate conformational changes of the Gi1 heterotrimer during GTP-induced activation (Figure 3a). This begins with nucleotide binding, reflected in the GTP-Primed state, wherein the phosphates of GTP become coordinated by hydrogen bonding with the backbone nitrogens of α1 residues E43 to T48, resulting in a shift of interactions in the rest of the α1 helix and the rerouting of the top of α1-linker I region (K54 to S62, Figure 3b, Extended Data Figure 3b). The linker rerouting facilitates its substantial ordering, which in turn leads to the ordering of the AHD in the open state. Next, we observe the initial closure of the AHD in the G-ACT-1 state, where the AHD is positioned underneath the RHD but without full ordering of Linker I and the top of helix αA (Figure 3a,c). This is followed by a further shift of the AHD towards the RHD and nucleotide, and the complete ordering of Linker I and αA helix in the G-ACT-2/3 states (Figure 3c).

The C-terminal α5 of the G-ACT-1 complex remains engaged with the receptor, but the map density in that region is modest and makes registering the helix challenging, reflecting in part its conformational heterogeneity. 3DVA analysis on this particle set produces maps with varying degrees of α5 helix engagement to MOR (Extended Data Figure 5a), similar to what was observed in our prior work with β_2_AR-Gs. Nevertheless, the locally refined map for the G-protein is sufficient to model backbone and bulky side chains, including around the α5 region. When the G-protein transitions from the GTP-Primed state to G-ACT-1 we observe significant shifts in α1. This includes the introduction of H57 as the top of helix folds and the entire α1 shifts away from α5, removing sidechain N52 from its position underneath the TCAT loop (T324-T327), a critical element for nucleotide base recognition (Figure 3d). In addition, we observe a small motion of the β2/3 loop and F191 away from α5, while H322 flips rotamers to bypass F334 and face towards the detergent micelle. Each of these changes would seem to clear steric blocks around α5 to allow for the substantial retraction and quarter-turn rotation that will occur when this helix fully pulls away from the receptor. This is consistent with mutagenesis studies showing that F334 and H322 play key roles in stabilizing the complex between rhodopsin and Gi^21^. Once the α5 is fully retracted in the G-ACT-2/3 states, we observe that the β2-β3 loop and F191 shift slightly back towards α5, with new hydrophobic packing forming in the F334-H322 and F336-H57-F191 regions (Figure 3e). These changes likely help stabilize the retracted α5 conformation and provide an energy barrier to re-insertion of the helix into the receptor. In support of this notion, experiments examining GDP release have also revealed that the bulky F336 sidechain of the α5 helix is necessary for preventing high basal activity by Gi1^22^.

As the AHD starts to close and gradually order while transitioning through all the G-protein states identified in this work, the switch I region (also known as linker II, T177-G183) folds towards the nucleotide (Figure 3f). This includes T181 growing increasingly close to the site of the bound catalytic magnesium ion, in concert with changes in D200 of switch II (V201-F215) which will also form part of the magnesium coordination shell. In the switch regions (I-III), which help control the interaction between Gα and Gβγ, we observe that the initial closure of the AHD in G-ACT-1 starts to bring switch III (A232-D242) towards switch II, with even closer proximity when the AHD orders in G-ACT-2 (Figure 3h). Switch II, in contrast, does not change its conformation until the AHD fully orders, driving a slight rearrangement of its coil portion towards the bound nucleotide.

In the case of DAMGO and MP, 3DVA identifies an additional state, G-ACT-3, where we observe the switch II region moving away from Gβγ (Supplementary Video 2,3). This state shows substantial rearrangement of several hydrophobic residues including W258 and W207, generating an increase in internal hydrophobic packing of the Gα as the entire subunit pulls away from Gβγ (Figure 3g,i). This is accompanied by an attraction between R205 and the polyanionic portion of switch III, and R208 forming a salt bridge with E245. The seemingly central role of these residues in this conformational change is consistent with mutagenesis studies that have identified R208 and E245 as two out of three residues (alongside G203) forming the ‘triad’ required for GTP-induced G-protein activation^23^. Finally, in the G-ACT-3 state, the coil portion of switch II (V201-E207) pulls in towards the GTP to position G202 and G203 closer to the nucleotide phosphates (Extended Data 5b). Notably, this situation is contrasted with LFT, where 3DVA analysis of the G-protein in G-ACT-2/3 reveals a different array of states (Supplementary Video 4). The first is virtually identical to the G-ACT-2 state of DAMGO and MP where Gα and Gβγ are close, with the exception of a weakening in density for the switch II region containing G202 and G203 (Extended Data Figure 5c, Supplementary Video 4). In the second state revealed by 3DVA, Gα has separated from Gβγ to a lesser extent than with MP and DAMGO, while the base of switch II has started to move towards Gα without fully interacting with the RHD (Extended Data Figure 5d,5e, Supplementary Video 4). It is possible that the presence of favorable interactions found in MP and to a lesser extent DAMGO between the stable, inactive-like receptor and the G-protein heterotrimer stabilizes the G-ACT-3 state and allows it to be observed in our cryo-EM experiments. On the other hand, we postulate that the lack of LFT-bound receptor-Gi interactions allows the G-protein to continue from this heterotrimer separated state to rapid dissociation of Gα and Gβγ, as also observed in our experiments.

To probe this hypothesis, we performed MD simulations of the G-ACT-3 state with either G-protein alone, the MP G-ACT-3 complex obtained here, or a state constructed from aligning an LFT-bound active-state receptor to its equivalent in MP-bound G-ACT-3 (Figure 4). In the simulations of G-protein alone, 40% of the trajectories show that the αN helix, one of the last major points of connection between Gα and Gβγ, begins to unfold (Figure 4a). This contrasts with our simulations of the MP-bound G-ACT-3 complex, which remain largely stable in the cryo-EM-resolved structure without any sustained unfolding of the αN helix (a single trajectory has a minor unfolding event followed by refolding). This is supportive of the notion that the inactive-like receptor in MP-bound G-ACT-3 is serving as a kinetic ‘trap’ to help stabilize this arrangement of the G-protein. Also, while there’s no unraveling of the αN helix in MD simulations of the LFT-bound G-ACT-3-like complex, we observe poor engagement of the receptor, consistent with our cryo-EM results, that would likely lead to the complex dissociation. This is particularly prominent at the interface between MOR TM5-TM6 and the Gα C-terminus, which remains strongly engaged throughout the MP trajectories averaging ∼55% of the contacts observed in the structure (Figure 4b, Extended Data Figure 4e). In contrast, few of the LFT-bound trajectories ever reach this extent of G-protein C-terminus engagement, even though several LFT-bound trajectories have equivalent levels of ICL3 interaction with the cleft between Gα and Gβγ (Figure 4c, Extended Data Figure 4e). This suggests that perhaps similar to Mb6, which assumes a similar binding pose to the Gα C-terminus but binds only to the inactive state of GPCRs, the interaction between the Gα C-terminus and the outside of the TM5-TM6 segment is receptor state-dependent.

**Figure 4:**
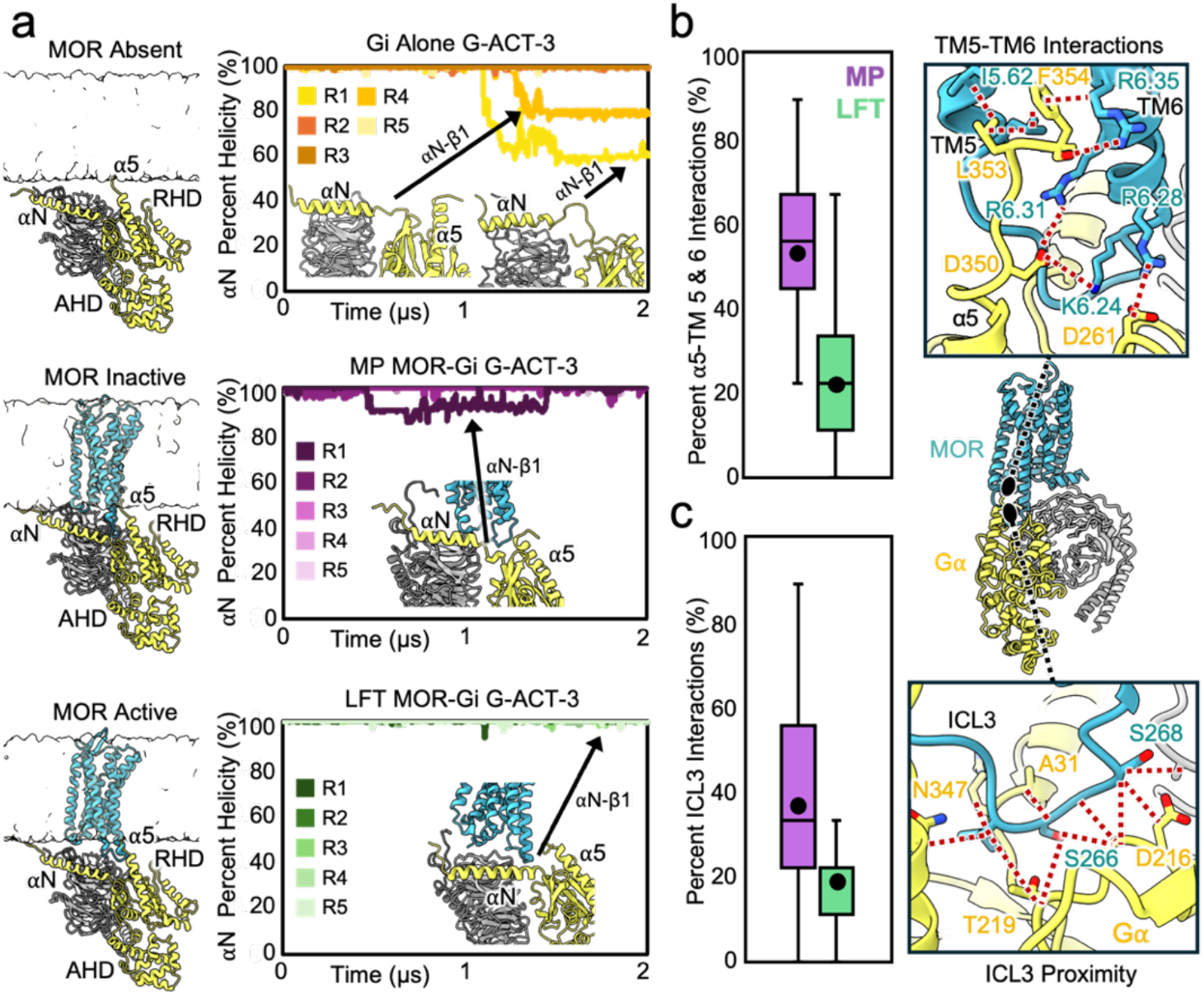
MD Simulations Probing the Stability of G-ACT-3. **a)** Simulation setups and percent helicity plots of the αN helix of Gαi over 2 μs MD simulations of the G-ACT-3 Gi without receptor, with the MP-bound inactive receptor, and with a LFT-bound active receptor. **b)** Diagram of the interactions between TM5-TM6 of MOR and the C-terminus of Gαi in the G-ACT-3 MP structure and box-and-whisker plot of the percent of these interactions maintained over 2 μs MD simulations. The center line corresponds to the median, box limits upper and lower quartiles, whiskers 1.5x interquartile range, and the black circle denotes the mean. **c)** Diagram of the contacts between ICL3 and the Gα/μγ cleft in the G-ACT-3 MP structure and box-and-whisker plot of the percent of these interactions maintained over 2 μs MD simulations. The center line corresponds to the median, box limits upper and lower quartiles, whiskers 1.5x interquartile range, and the black circle denotes the mean.

### Divergent Gi1 and Gs Activation Dynamics

To our knowledge, there are few GPCRs reported to strongly and primarily couple to both the Gi and Gs subfamily of G-proteins, presumably owing to their opposing actions on adenylyl cyclase (inhibitory vs. stimulatory). Thus, juxtaposing the behavior of Gi and Gs activation between canonical, strong-coupling receptors, such as the MOR and β_2_AR, is relevant and allows for potential comparisons in endogenous signaling profiles. In support of this, DEER studies of RHD-AHD distances for the rhodopsin-Gi complex showed very similar behavior to what we observe for MOR-Gi(17), suggesting that our observations on Gi vs Gs differences may be generalizable to other GPCRs. Comparison of the results of this cryo-EM study with MOR-Gi to our prior study with β_2_AR-Gs points to mechanistic differences between the two G-proteins. A striking deviation occurs in the baseline dynamics, wherein the nucleotide-free MOR-Gi does not display AHD closure, while β_2_AR-Gs displays intrinsic opening and closing of AHD (Figure 4a). Binding of GTP to MOR-Gi drives a substantial ordering of the linker region and rigidification of the AHD, encouraging its closing against the RHD (Figure 4a). This indicates that an induced conformational change drives AHD closure in MOR-Gi, in contrast to β_2_AR-Gs that appears to leverage conformational capture of AHD when nucleotide is present (Figure 5a).

The difference in the mechanism of AHD closure between MOR-Gi and β_2_AR-Gs appears to stem from the divergence between the sequences and behavior of the linker I and αA helix regions of the two G-proteins (Figure 5b,c). The linker itself bears a phenylalanine in place of Y61 in Gi, and is one amino acid longer for Gs, terminating in a glycine residue that is absent for Gi. On the αA helix, Gi has several tyrosine residues, including Y69, that are substituted by lysines in Gs. Both Y61 and Y69 make hydrogen bonding interactions with Gβγ in the open ordered state of GTP-bound Gi (ED Fig. 3d) and pack hydrophobically against the AHD in the closed GTP-bound conformation (Figure 5b). The amino acid differences in Gs result in αA tilting much further away from the rest of the AHD in the GTP-bound closed conformation while much of the top of αA and linker I are disordered, traits shared with crystal structures of Gs-GTP^24^ with the AHD in closed conformation (Extended Data Figure 6a).

**Figure 5:**
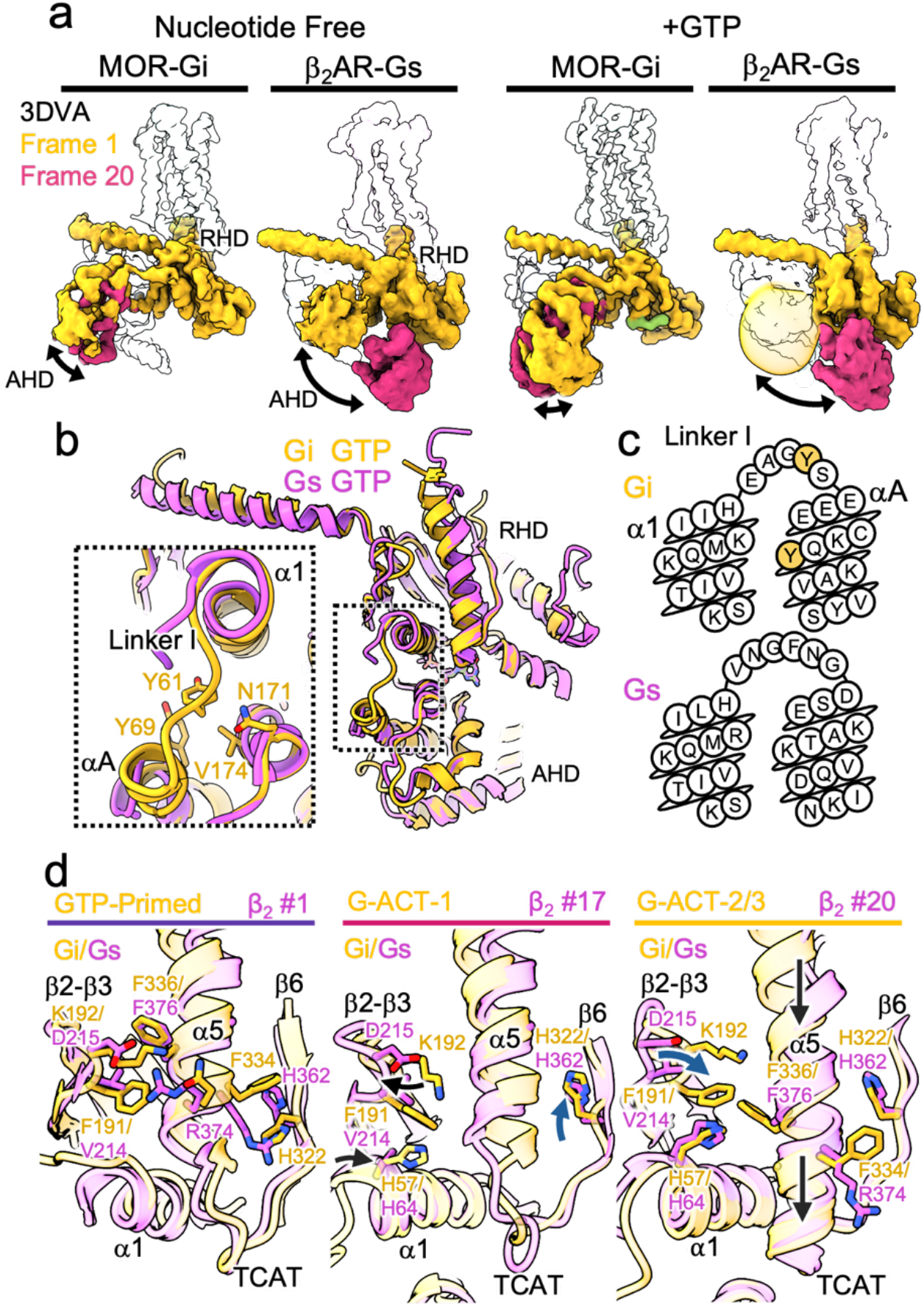
Differences in Gi vs Gs Activation Dynamics. **a)** Comparison of the 3DVA analysis results for the first principal component of the nucleotide free (left) and GTP-bound, predominant particle stack state (right) between MOR (teal)-Gi and β_2_AR (purple)-Gs. **b)** Alignment of Gi (yellow) and Gs (pink) in an AHD closed, GTP-bound state (G-ACT-2 for MOR-Gi, 3DVA frame 20 for β_2_AR-Gs). The inset is centered on the linker I region, highlighting the packing of αA and linker I against the rest of the AHD in Gi that is absent in Gs. **c)** Comparison of the sequences of the α1-linker I-αA sequences of Gi1 and Gs(short). **d)** Comparison of Gi (yellow) and Gs (pink) from roughly equivalent states in their cryo-EM trajectories for GTP-induced activation from their respective receptors.

### Agonist-Dependent Free Energy Landscape of GTP-Induced MOR-Gi Activation

Together these results provide a qualitative view of the free energy landscape of MOR-Gi activation by GTP and how different agonists shift this landscape to impart their distinct signaling profile (Figure 5a). Based on the present studies and DEER work with Gi^17^, in the absence of nucleotide, the energy of the open AHD state would seem to be substantially lower than that of the closed AHD state, with a sizable barrier between them. Binding of GTP induces shifts in the α1 helix, producing an ordering of the linker and AHD and reducing the relative energy of the closed AHD state relative to the open state, thus enabling AHD closure (Figure 6, Extended Data Figure 7a). The closure generates contacts between switch I and switch II, the AHD and switch III-α3, and the AHD and RHD, driven in part by new contacts between GTP and the AHD (Extended Data Figure 7a). In this AHD closed state (G-ACT-1), the α5 helix has minimally shifted its contacts with the receptor, and the linker I-αA remains highly dynamic. These interactions seemingly prime the complex to transition to full closure and ordering of linker I-αA, which is observed in the G-ACT-2 state along with full retraction of the α5 helix and downward shift of the TCAT loop to more extensively contact the nucleotide (Extended Data Figure 7a).

**Figure 6:**
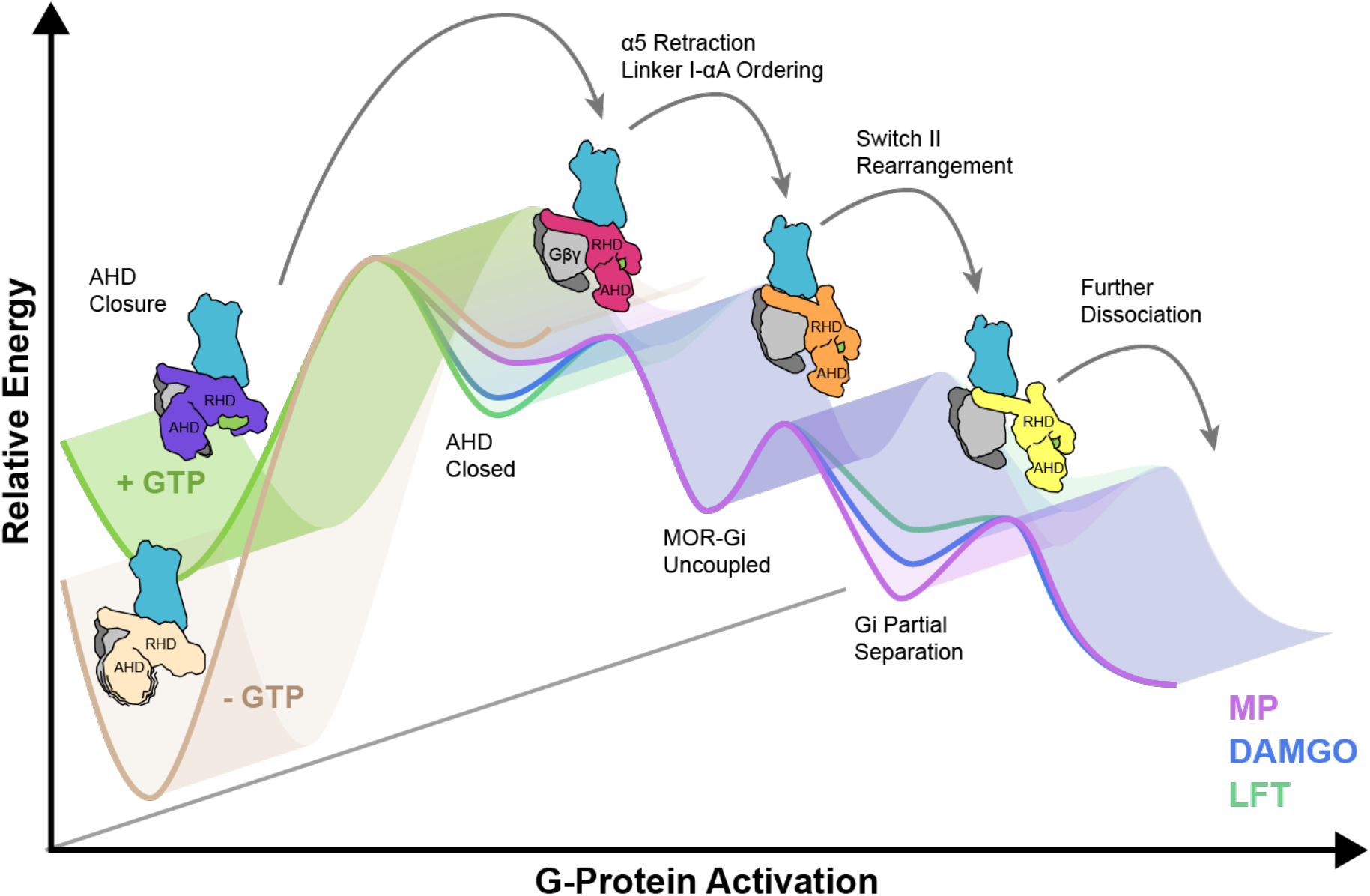
Proposed Ligand-Dependent Free Energy Landscape of GTP-Induced MOR-Gi Activation. Cartoon representation of the free energy landscape proposed in this work. Addition of GTP shifts the relative free energy of the AHD open and closed states by ordering the linker region between the RHD and AHD, and triggering a cascade of conformational changes towards G-protein activation and dissociation. Partial agonists appear to stabilize a potential kinetic ‘trap’ state near the end of the process, with partial separation between Gα and Gβγ.

According to our results, and in alignment with recently published DEER measurements of MOR TM6 conformations^9^, the receptor occupies a ligand-dependent ensemble of states, with weaker agonists tending towards inactive-like states while stronger agonists producing more dynamic receptor states. The fully closed AHD generates further contacts with switch III and α3, pushing these regions closer to switch II. This appears to prime the conformation of Gα for the extensive switch III/α3 and switch II contacts needed to stabilize the pulling of switch II away from Gβγ and towards the nucleotide-binding pocket (Extended Data Figure 7a). The inactive-like receptor state allowed by weaker agonists is captured in the pseudo-stable complex with the G-protein heterotrimer of MP-bound G-ACT-3. Those complexes display a more separated Gα from Gβγ where the switch II and switch III-α3 contacts have formed and the RHD of Gα has started to pull away from Gβγ (Figure 6, Extended Data Figure 7a). In this way, weaker agonists may stabilize a state that serves as a ‘kinetic trap’ along the GTP-induced activation pathway, preventing full dissociation of the G-protein subunits from each other and the receptor itself. This is supported by the MD simulations of the G-ACT-3 state in the presence and absence of MOR, in which Gi αN unfolding and separation from Gβγ only occurs without the MOR present. Further, while we could resolve the G-ACT-2 state and heterogeneity in the switch II region of the LFT data, where the receptor seems to remain in a dynamic, primarily active-like state, we could not resolve a stable G-ACT-3 state, which we attribute to rapid further dissociation of the G protein heterotrimer.

The findings point to at least two possible rate-limiting steps. The first potentially high barrier step in G-protein activation upon GTP loading is the closure of the AHD, followed by progression to states of increasingly tighter closure against the RHD-GTP module with the beginning of α5 retraction from the receptor core and destabilization of ICL2 and ICL3 interactions. In our experimental framework, it is likely that in the case of LFT and DAMGO, slow AHD closure is followed by rapid progression through the remaining uncoupling steps. Nevertheless, we also cannot rule out that steps of the GTP-induced dissociation are reversible and there likely exists some complex transitioning backward along the canonical GTP activation pathway. The second rate-limiting step is the full withdrawal of the α5 succeeded by further complex dissociation. This is underlined by a notable difference in Gi and Gs that we observe in the α5 helix region. When the G-protein is still fully engaged with the receptor, both Gαi and Gαs form substantial internal interactions involving the β2-β3 loop, α1, and α5. These interactions are common despite differences in β2-β3 loop composition (F191 and K192 for Gi vs. V214 and D215 in Gs, Figure 5d). For α5 to retract and disengage from the receptor, H322 of Gi will need to flip rotamers past F334. In contrast, H362 of Gs, which is already flipped upward, does not present the same steric barrier to the α5 helix R374 (Figure 5d). In the intermediate state(s) of Gi and Gs activation, the β2-β3 loop and α1 pull away from α5 in both G-proteins (Figure 4d). In Gs, however, the backbone of the β2-β3 loop continues to move further from α5, as also observed in crystal structures of Gαs bound to a GTP-analogue^24^ (Extended Data Figure 6b). This contrasts with Gi, where the β2-β3 loop pulls back towards the α5 helix, forming hydrophobic packing with F336, a key residue for allosteric communication with the nucleotide-binding site^22^. Together these results demonstrate distinct differences in both the number and the character of interactions between α5 and the rest of the RHD which provide barriers for the extension/retraction of α5 between the inhibitory Gi and the stimulatory Gs. This is also consistent with recent studies demonstrating that interactions between the α5 and the β2-β3 loop are important for driving G-protein subtype selectivity by affecting GPCR-G-protein activation kinetics^25^. For both potential rate-limiting steps, AHD closure and full α5 withdrawal, it would appear that Gi faces higher energy barriers compared to Gs, as we observe the majority of GTP-loaded complexes with an open-state ordered AHD without changes in receptor Gi protein interactions. This behavior may be partly behind the slower GTP exchange rate observed for Gi isoforms compared to Gs^26-28^.

## Discussion

Here we show how building upon cryo-EM studies of static receptor-transducer snapshots toward dynamic ensembles of structures under non-equilibrium conditions can help address unresolved questions of GPCR signaling mechanisms. This work has revealed significant diversity in the GTP-induced activation pathway, both between two distinct G-proteins, but also for the same receptors bound to different ligands. We provide structural evidence of how ligands of distinct efficacies affect not only receptor conformations but aspects of the GPCR-G-protein interactions during the signaling process. We have identified a ligand’s promotion, or lack thereof, of a persistently dynamic and open conformation of TM6 as key for modulating the GTP-induced activation of Gi by MOR. Further, our findings suggest that partial agonists allow MOR to easily revert to a TM6-inward conformation which may form transient, pseudo-stable complexes with activated G-proteins, stalling downstream signaling. We note that the rapidly adopted inward conformation of TM6 also explains the relatively weak propensity of partial agonists to promote arrestin and also potentially GRK recruitment, a characteristic recently described for MOR^6,14^ but likely involving several other GPCRs. Given these results, we anticipate non-equilibrium and time-resolved cryo-EM will likely reveal that there is significantly more structural, kinetic, and dynamic diversity in the landscape of GPCR-G-protein interaction process than previously revealed by nucleotide-depleted structures.

We also note several caveats to our experimental approach. First, given the limitation in temporal resolution to the level of seconds and the complexes seemingly reaching pseudo-steady state within the window of our experiment, the cryo-EM data alone cannot fully report on the reversibility of the proposed processes or the specific transition pathways. While our complementary MD simulations have helped to fill in some of the gaps, there may very well be ‘backwards’ transitions in the process in our sample. Further, our study is performed with purified components in vitro, in the absence of the native lipid bilayer and any other endogenous proteins that may affect G-protein signaling, which may substantially modify the free energy landscape of this process *in vivo*. Finally, we have examined only the second half of the GPCR-G-protein signaling, i.e. complex dissociation upon GTP addition, and not G-protein association with the receptor. Further time-resolved cryo-EM studies of the GDP-bound G-protein interacting with the receptor to trigger AHD opening and nucleotide release will likely be equally as informative and valuable for understanding ligand pharmacology.

## Supporting information

All Extended Data Figures

## Acknowledgements

St Jude GPCR collaborative grant (G.S.), NIH R01 DA059978 (S.M.), NIH K99HL16140601 (M.M.P.-S.), NIH K99/R00 HD107581 (M.J.R.), and CPRIT award RR230042 (M.J.R.). M.J.R. is a CPRIT Scholar in Cancer Research. We thank Nevin Lambert, Roger Sunahara, and Jonathan Javitch for discussions and critical reading of the manuscript.

## Contributions

M.J.R. expressed and purified proteins, prepared complex, prepared cryo-EM grids, collected, analyzed and processed cryo-EM data, built and refined atomic models, designed and executed molecular dynamics simulations, analyzed data, and prepared figures. M.M.P.-S. collected data for MP timepoints. M.C.P. expressed and purified G-proteins. B.R.V. synthesized compounds under supervision of S.M.. G.S. conceived and supervised the project. M.J.R. and G.S. wrote the manuscript.

## Data availability

The atomic coordinates of the LFT-bound states have been deposited in the Protein Data Bank (PDB) under accession codes 9ODE, 9ODF, 9ODG, 9ODH, 9ODI and all associated maps in the Electron Microscopy Data Bank (EMDB) under accession codes EMD-70356, EMD-70357, EMD-70358, EMD-70359, EMD-70360, EMD-70361, EMD-70362, and EMD-70363. The atomic coordinates of the MP-bound states have been deposited in the PDB under accession codes 9ODJ, 9ODK, and 9ODL, with maps deposited in EMDB EMD-70364, EMD-70365, and EMD-70366. The atomic coordinates of the DAMGO-bound states have been deposited in the PDB under accession codes 9ODM, 9ODN, 9ODO, 9ODP with the associated maps in the EMDB under accession codes EMD-70369, EMD-70370, EMD-70371, EMD-70372, EMD-70373, and EMD-70374.

## Methods

### Expression and Purification of Gi1 Heterotrimer

Gi Heterotrimer was expressed in Sf9 cells with the baculovirus expression system as described previously^11^. Cell pellets were resuspended in a lysis buffer of 20 mM HEPES pH 7.5, 1 mM EDTA pH 8.0, 1mM MgCl_2_, 5% glycerol, 5 uM β-mercaptoethanol, 100 uM GDP, protease inhibitor cocktail, and benzonase. The lysate was stirred at 100 rpm for 20 minutes at 4C before 30 minutes of ultracentrifugation at 100,000xG. Membranes were dounce homogenized in a buffer of 100 mM NaCl, 20 mM HEPES pH 7.5, 5% glycerol, 1 mM MgCl_2_, 5 mM β-mercaptoethanol, 100 μM GDP, 1 mM benzamidine, protease inhibitor cocktail, benzonase, and 1% cholate. This solution was stirred for 60 minutes at 100 rpm at 4C before ultracentrifugation at 100,000xG for 30 minutes at 4C. Ni-NTA beads and 30 mM imidazole were added to solubilized G-protein and the solution was incubated for 1 hour. Beads were loaded onto a gravity column and washed with 10 CV of buffer containing 100 mM NaCl, 20 mM HEPES pH7.5, 5% glycerol, 1 mM MgCl_2_, 5 mM β-mercaptoethanol, 100 μM GDP, 30 mM imidazole, and with increasing concentrations of LMNG/CHS and decreasing concentrations of cholate until a final detergent concentration of 0.05% LMNG/0.005% CHS is achieved. The same buffer supplemented with 250 mM imidazole was used to elute G-protein and 1 mg HRV-3C protease/50 mg G-protein was added to cleave the 6xHis tag overnight dialyzing against low imidazole buffer. The following day the protein is flowed through a Ni-NTA column, washed, and both the flow through and wash were concentrated. Gi1 heterotrimer was then injected onto an ENrich 650 column in a buffer of 100 mM NaCl, 20 mM HEPES pH 7.5, 5% glycerol, 1 mM MgCl_2_, 100 μM TCEP, 20 μM GDP, 0.01% LMNG, 0.001% CHS. Gi1 heterotrimer was concentrated to 5-10 mg/ml and flash frozen in liquid nitrogen.

### Expression and Purification of MOR

MOR was expressed in Sf9 cells with the baculovirus expression system as described previously^11^. Pellets of MOR were thawed and suspended in a hypotonic lysis buffer containing 20 mM HEPES pH 7.5, 1 mM EDTA pH 8.0, 1 mM MgCl_2_, 100 μM TCEP, 10 μM Naloxone, protease inhibitor cocktail, and benzonase. Solution was stirred at 100 rpm at 4C for an hour before centrifugation at 100,000xG. Membrane pellets were then resuspended in 500 mM NaCl, 20 mM HEPES pH 7.5, 1 mM MgCl_2_, 10% glycerol, 10 μM Naloxone, 100 μM TCEP, 1 mM benzamidine, protease inhibitor cocktail, and 5% LMNG/1% CHS/1% cholate detergent stock solution was slowly added while stirring at 100 rpm at 4C to a final concentration of 1% LMNG, 0.2% CHS, 0.2% cholate. Solubilization proceeded for 3 hours, with 2 mg/ml iodoacetamide added at the 30 minute and 1 hour points. Solution was ultracentrifuged at 100,000xG to remove, the supernatant was supplemented with 20 mM imidazole, and the solution was gravity-loaded over Ni-NTA resin. The column was washed with buffer containing 500 mM NaCl, 20 mM HEPES pH 7.5, 20 mM imidazole, 0.1% LMNG, 0.01% CHS, 10 μM Naloxone, 100 μM TCEP; and eluted with a buffer containing 250 mM NaCl, 20 mM HEPES pH 7.5, 250 mM imidazole, 0.05% LMNG, 0.005% CHS, 10 μM Naloxone, and 10% glycerol. The MOR was supplemented with 5 mM CaCl_2_ and loaded onto M1 flag resin, washed with buffer containing 20 mM HEPES pH 7.5, 250 mM NaCl, 2 mM CaCl_2_, 0.01% LMNG, 0.001% CHS, 10 μM Naloxone; and eluted with buffer containing 20 mM HEPES pH 7.5, 250 mM NaCl, 1 mM EDTA, 0.01% LMNG, 0.001% CHS, 10 μM Naloxone, 10% glycerol, and 0.1 mg/ml FLAG peptide. The eluent was supplemented with 100 μM TCEP, concentrated, and loaded onto an ENrich 650 column for SEC in buffer containing 100 mM NaCl, 20 mM HEPES pH 7.5, 0.01% LMNG, 0.001% CHS, and 100 μM TCEP. Fractions containing monomeric MOR were concentrated and used immediately for G-protein complexation.

### Formation and Purification of MOR-Gi Complex

Purified apo MOR was incubated with 200 μM of either LFT, DAMGO, or E-MP for 1 hour on ice before the addition of a molar excess of purified G-protein heterotrimer. Complexation was incubated for 1 hour on ice before the addition of 2 μL apyrase solution. This reaction was allowed to proceed overnight on ice. The reaction solution was then diluted with buffer containing 100 mM NaCl, 20 mM HEPES pH 7.5, 0.01% LMNG, 0.001% CHS, 5 mM CaCl_2_, and 20-100 μM agonist; then loaded over M1 FLAG resin. Resin was washed with buffer containing 100 mM NaCl, 20 mM HEPES pH 7.5, 0.005% LMNG, 0.0005% CHS, 1 mM CaCl_2_, and 10-50 μM agonist, and eluted into a buffer containing 100 mM NaCl, 20 mM HEPES pH 7.5, 0.002% LMNG, 0.0002% CHS, 1 EDTA, 0.1 mg/ml FLAG peptide, 10% glycerol, and 20-100 μM agonist. Once eluted, 100 μM TCEP was added and complex was concentrated and then loaded onto a SEC Enrich 650 column in buffer of 100 mM NaCl, 20 mM HEPES pH 7.5, 0.001% LMNG, 0.0001% CHS, 0.00033% GDN, 100 μM TCEP, 2 mM MgCl_2_, and 1-100 μM agonist. Complexes were then concentrated to 15 mg/ml for cryo-EM.

### Cryo-EM Sample Preparation & Data Collection

All samples were prepared on glow-discharged holey carbon grids with gold support (Quantifoil R1.2/1.3). 3 μL of MOR-Gi sample was loaded into an FEI Vitrobot Mark IV with the chamber held at 4°C and 100% humidity. 0.3 μL of 10 mM GTP was pipetted onto the grid concomitantly with pressing the foot pedal of the Vitrobot, with blot time set to 3 seconds and wait time set to produce the desired total process time until plunging in liquid ethane, noting that the vitrification process takes an additional 3 seconds in addition to the blot and wait time. Cryo-EM data sets were collected on a Titan Krios electron microscope at an accelerating voltage of 300 kV with the SerialEM software^28^ and a Gatan K3 Direct Electron Detector/Bioquantum energy filter for DAMGO or the EPU software and a Falcon 4i direct electron detector/Selectris energy filter for LFT and MP. Number of micrographs and electron doses for each dataset are provided in Extended Data Table 1, each entry represents an independent cryoEM grid.

### Data Processing

For datasets collected on the Falcon 4i, .eer files were converted to tiff format with the relion_image_handler utility. All tiff files were then loaded into RELION-4.0^29^ for motion correction with MotionCor2^30^, CTF estimation with CTFFIND4^31^, and template-based particle picking. Binned extracted particles were then imported into CryoSPARC^32^ for 2D and 3D classification. Particle stacks were then refined with the nonuniform refinement routine and transferred back to RELION for re-extraction of unbinned particles. For structures that were likely to refine below 3.0A resolution, particles were refined in RELION and subjected to Bayesian polishing^33^. All particle stacks were then imported back into CryoSPARC for final non-uniform refinement, local refinement, and/or 3DVA processing^16^.

### Model Building

Initial models were based off of our prior cryo-EM structures of MOR-Gi with DAMGO, MP, and LFT, crystal structures of GDP-bound Gi heterotrimer, and our inactive MOR cryo-EM structure (PDB:6DDE, 1GP2, 7T2G, 7T2H, 7UL4). Manual model building was performed in Coot^34^ with refinement in Phenix^35^. GlideEM^36^ was used to validate the ligand poses with the higher resolution maps as well as the poses for GTP. Details of cryo-EM map and model refinement are found in Extended Data Tables 2-4. Euler angle distribution plots, FSC curves, and selected map-model agreement panels are provided in Supplementary Information Figures 1-3.

### Molecular Dynamics Simulations & Analysis

In all systems, the N-terminus of Gα, the C-terminus of MOR, and the C-terminus of Gγ were built out to include their sites of lipidation (if not already modeled) by using residual low resolution map features for that respective state. All systems were oriented with the PPM webserver^37^ and then solvated in a box of POPC/CHS, TIP3P water^38^, 100 mM NaCl, and 10 mM MgCl2 (in simulations with GTP) using the CHARMM-GUI^39^. Solvated systems were simulated with the NAMD software^40^ with the CHARMM36 forcefield^41-43^ using a Langevin thermostat and Nose-Hoover Langevin piston barostat at 1 atm with a period of 150 fs and decay of 75 fs. Periodic boundary conditions were used with nonbonded interaction smoothing at 10 A to 12 A with long-range interactions handled with particle mesh Ewald. A 2 fs timestep was employed throughout the full simulations with SHAKE and SETTLE algorithms used. Harmonic restraints of 1 kcal/mol/Å^2^ were applied to all non-hydrogen, non-water/ion atoms and the systems were minimized for 1,500 steps before gradual heating from 0 to 303.15K in 20K increments with 0.4 ns of simulation per increment, with an additional 10 ns of equilibration at 303.15K. An additional 10 ns of equilibration with harmonic restraints only applied to non-hydrogen protein atoms, followed by another 10 ns of equilibration with harmonic restraints only applied to CA atoms. An additional 30 ns of unrestrained simulation was counted as equilibration, with the following simulation considered production. All simulations were run with 5 replicates with different initial seeds for random assignment of velocities. Systems were analyzed with VMD^44^ and python scripting. Helicity was calculated with dihedral angles falling into ϕ between 0° and -120° and φ between -90° and 15°. ICL3 interactions were calculated between MOR S268:α E216, MOR S268:α E216, MOR S268:α E216, MOR S268:α E216, MOR S268:α E216, MOR G267:α A31, MOR G267:α G217, MOR S266:α A31, MOR S266:α G217, MOR L265:α N347, MOR L265:α V37, MOR L265:α T219, MOR S268:β A56, MOR L265:α R32, and MOR S268:α E217. TM5 & TM6 interactions were calculated between the α C-terminal carbonyl:MOR R280, α F354:MOR R280, α L353:MOR I256, α F354:MOR I256, α D261:MOR R273, α D261:MOR K269, and α D350:MOR R276.

