## Supplementary material for "Non-Equilibrium Snapshots of Ligand Efficacy at the μ-Opioid Receptor": All Extended Data Figures

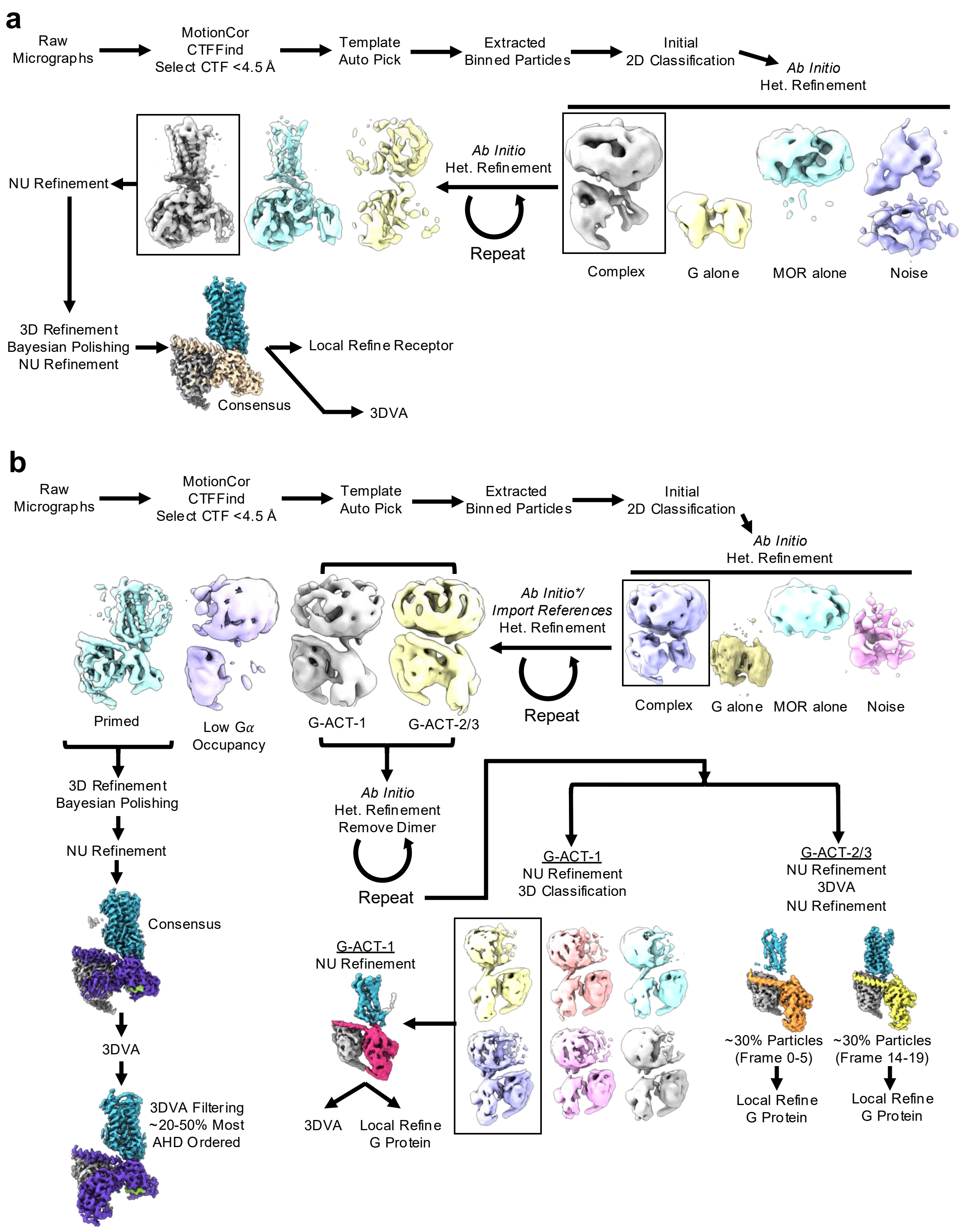


**Extended Data Figure 1: Cryo-EM data processing workflow a)** General processing workflow for the nucleotide-free datasets. **b)** General processing workflow for the GTP-bound datasets.

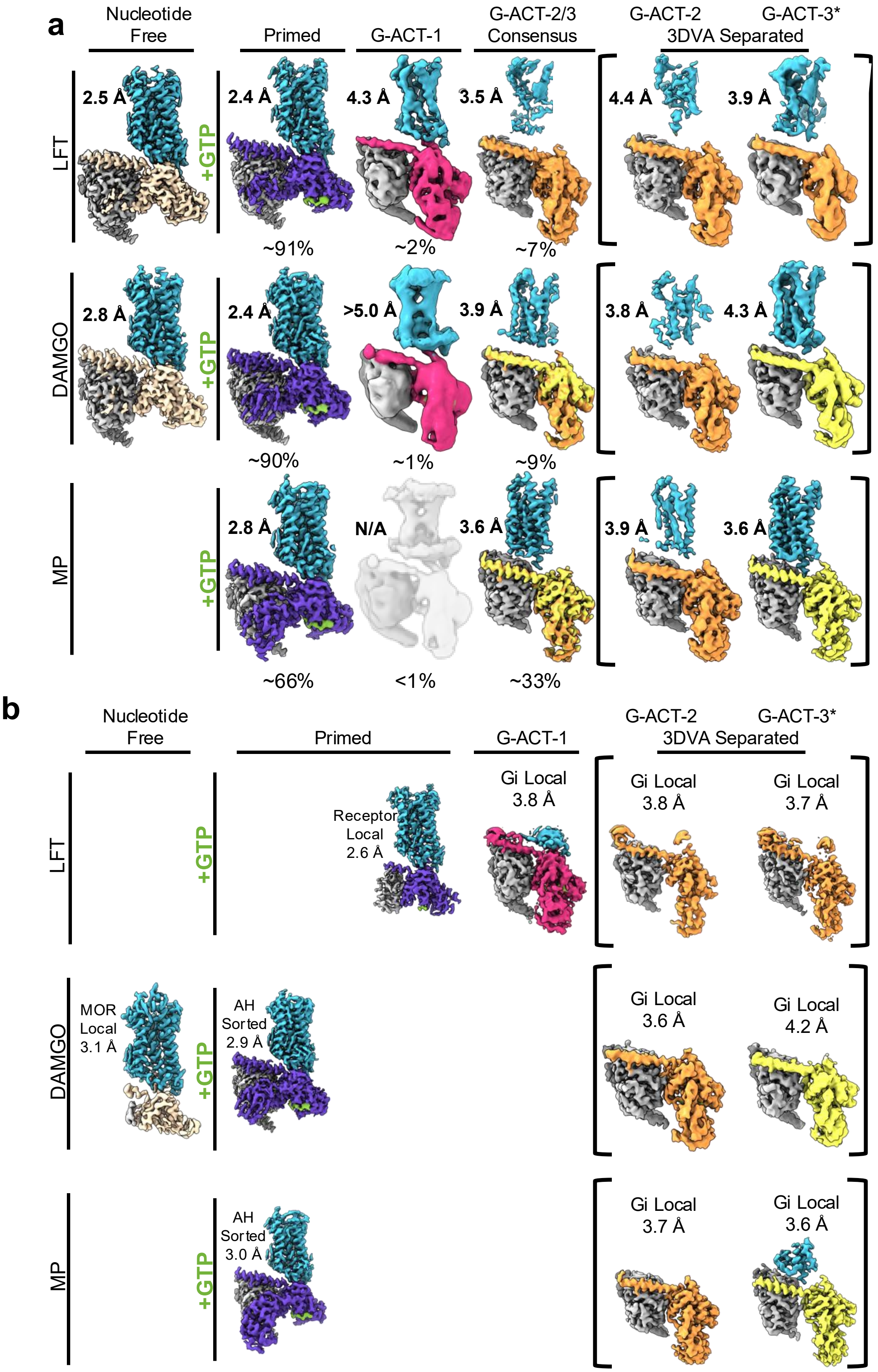


**Extended Data Figure 2: Cryo-EM maps of MOR-Gi bound to MP, DAMGO, and LFT in the baseline, G-Primed, G-ACT-1, G-ACT-2, and G-ACT-3 states. a)** Maps for the nucleotide free state, each of the consensus refinements of the three main ensembles of states for the GTP-bound state, and the 3DVA-separated states of the G-ACT-2/3 ensemble with each of the three ligands. G-ACT-3* corresponds to G-ACT-3 for MP and DAMGO, and G-ACT-2' for LFT. **b)** Additional maps generated in this work with local refinement and/or additional 3DVA/3D classification.

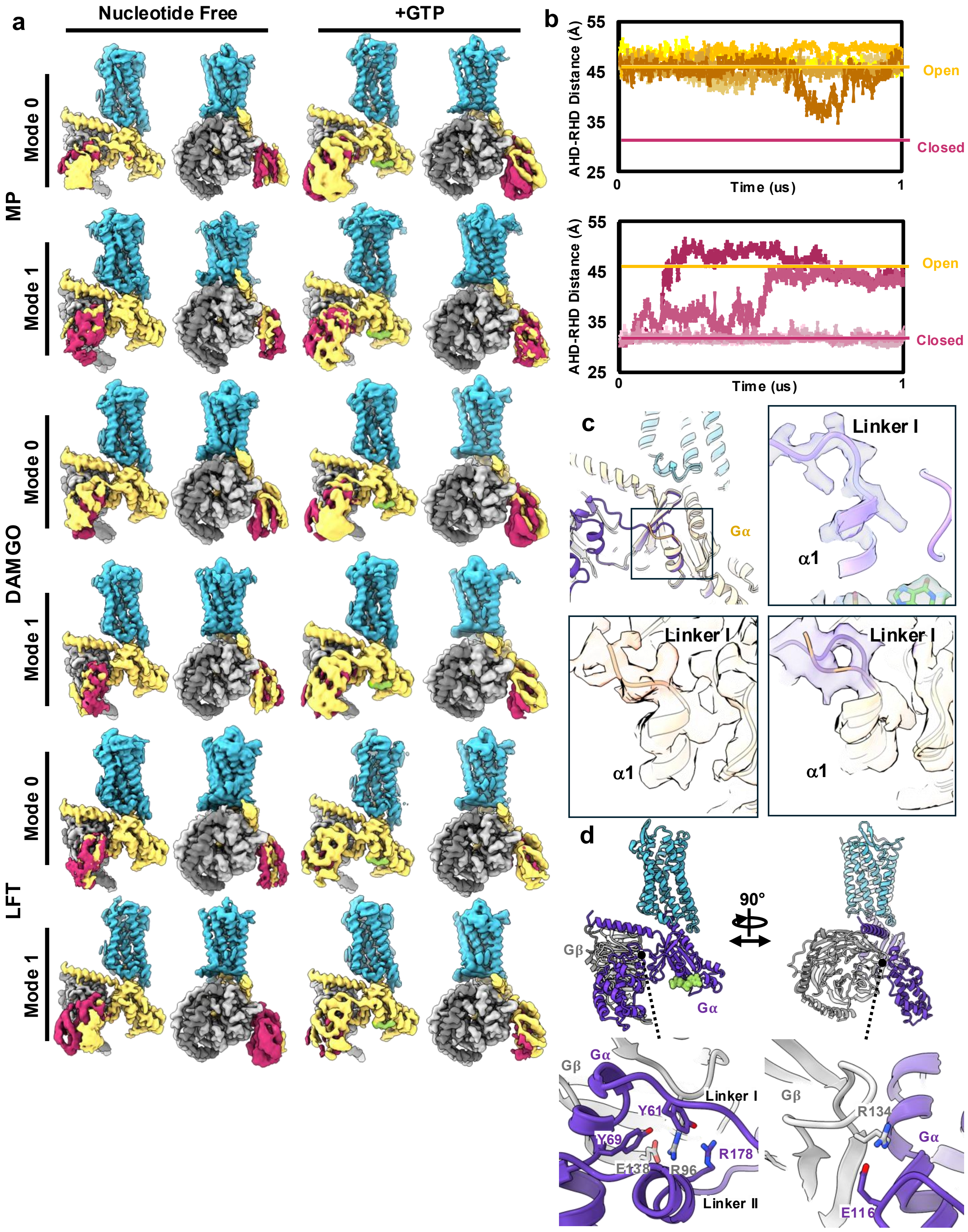


**Extended Data Figure 3: AHD motions in the nucleotide-free and GTP-bound Gi heterotrimer. a)** Comparison of the 3DVA analysis results for mode 0 and 1 of the nucleotide free (left) and G-Primed state (right) with each of the three ligands. **b)** AHD-RHD distances from nucleotide-free MD simulations where the AHD starts open (top, yellow gradient traces) or closed (bottom, magenta gradient traces). AHD-RHD distances from open and closed structures are drawn as straight lines. **c)** Comparison of the linker I region of Gi1 in the presence (purple) and absence (yellow-orange) of GTP. Maps are for the GTP bound state (top right), consensus nucleotide free (bottom left), and 3DVA-filtered for the presence of ordered linker (bottom right). **d)** Interactions between the AHD of Gα and Gβ in the GTP-bound, G-Primed state.

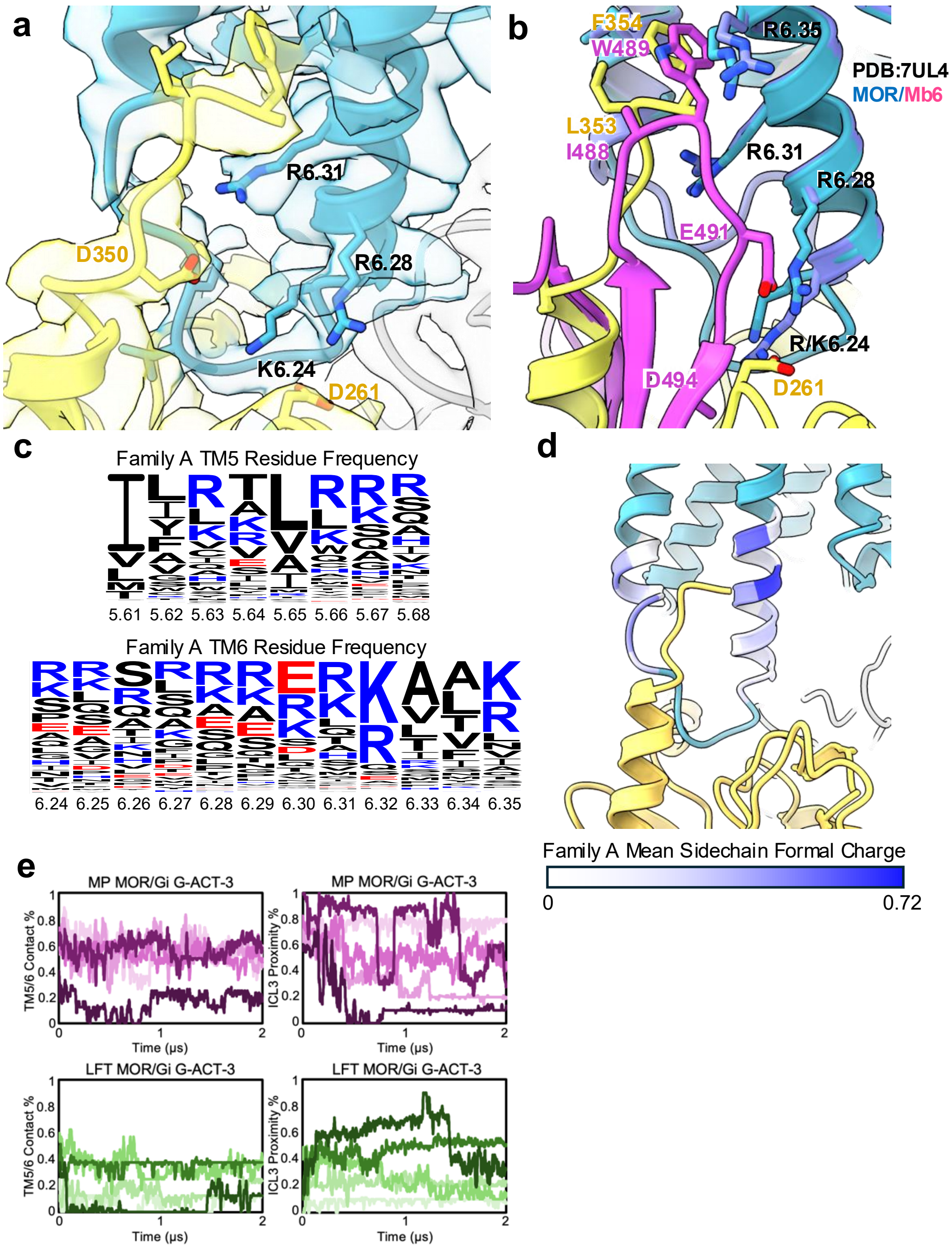


**Extended Data Figure 4: Interactions between Gα and MOR TM5-TM6 in the G-ACT-3 state a)** MP G-ACT-3 state model-map comparison highlighting the interface between TM5-ICL3-TM6 of MOR (aqua) and the C-terminus-RHD of Gα (yellow) **b)** Overlay of MP G-ACT-3 with the structure of MOR-Mb6 complex (Mb6 in magenta, PDB:7UL4). **c)** Residue frequency plot for the base of TM5 and TM6 across all human family A GPCRs. **d)** Analysis of the mean sidechain formal charge across all family A GPCRs at the base of TM5 and TM6. **e)** Time course of percent of TM5-TM6 Gi interactions and ICL3 proximity interactions in the G-ACT-3 simulations with MP (top, purple traces) and LFT (bottom, green traces) over 2 microsecond MD simulations. Lines plotted are rolling averages of 9 frames of MD simulation from five independent replicates.

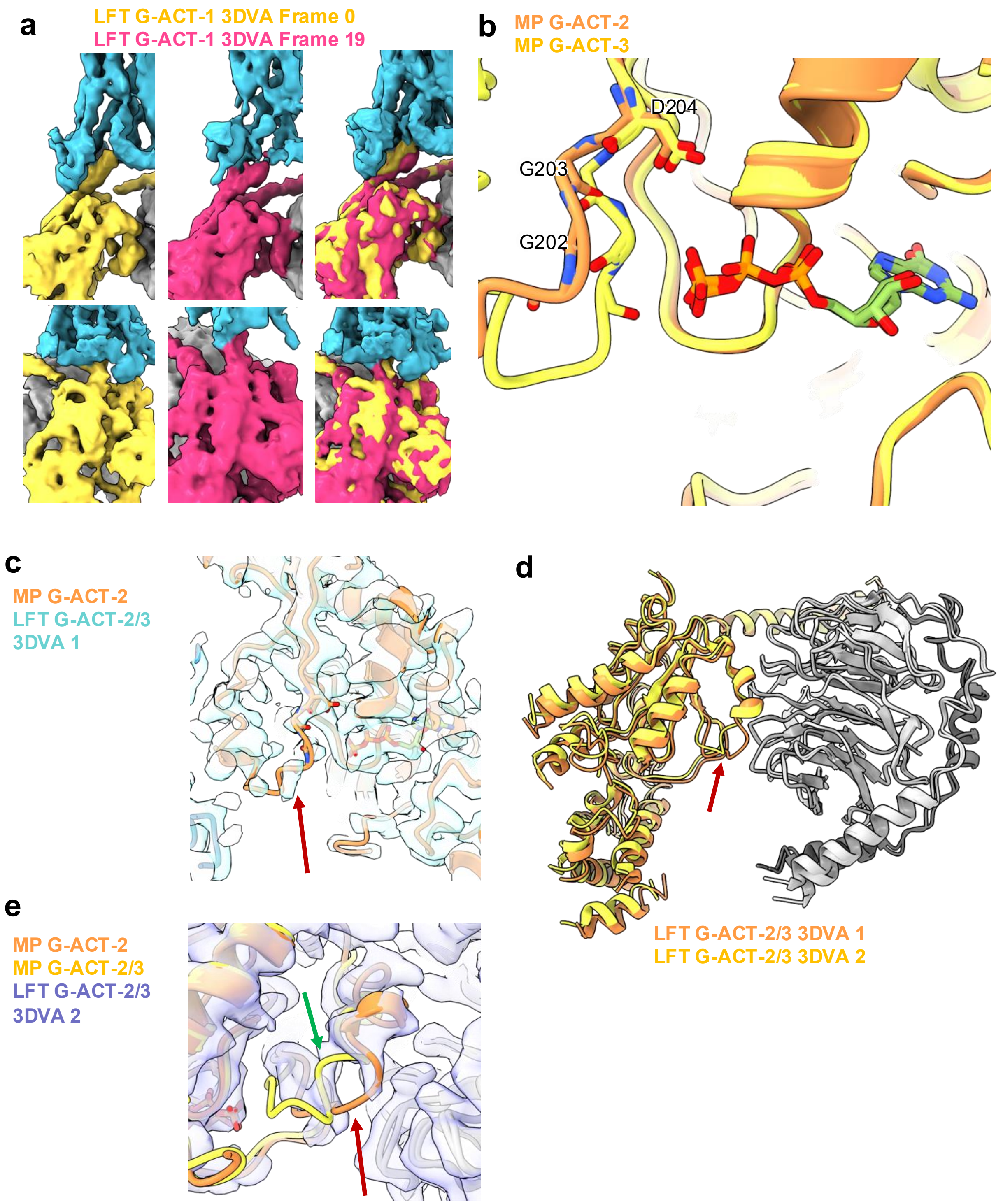


**Extended Data Figure 5: 3D Variability Analysis of the G-ACT-1 and G-ACT-2/3 states. a)** 3DVA results for LFT MOR-Gi G-ACT-1 state particles with Gα colored yellow for frame 0 and magenta for frame 19 of the principal component. **b)** Comparison of the coil portion of the switch II region for MP G-ACT-2 and G-ACT-3 highlighting the differences in G202 and G203. **c)** Comparison of the MP G-ACT-2 model with the map obtained from 3DVA filtering of the LFT G-ACT-2/3 particles, pooling the initial 1/3 of particles from the principal component. Arrow denotes the region of Switch II near GTP that is more disordered in the map **d)** Alignment on Gβγ of the structures modeled in the maps from LFT G-ACT-2/3 particles 3DVA analysis filtering, both the initial (3DVA 1) and final (3DVA 2) 1/3 of the particles. **e)** Comparison of the models for MP G-ACT-2, MP G-ACT-3, and the map obtained from 3DVA filtering of the LFT G-ACT-2/3 particles, pooling the final 1/3 of particles from the principal component. Green arrow highlights new map features and red arrow highlights where map features were lost compared to LFT G-ACT-2/3 3DVA 1.

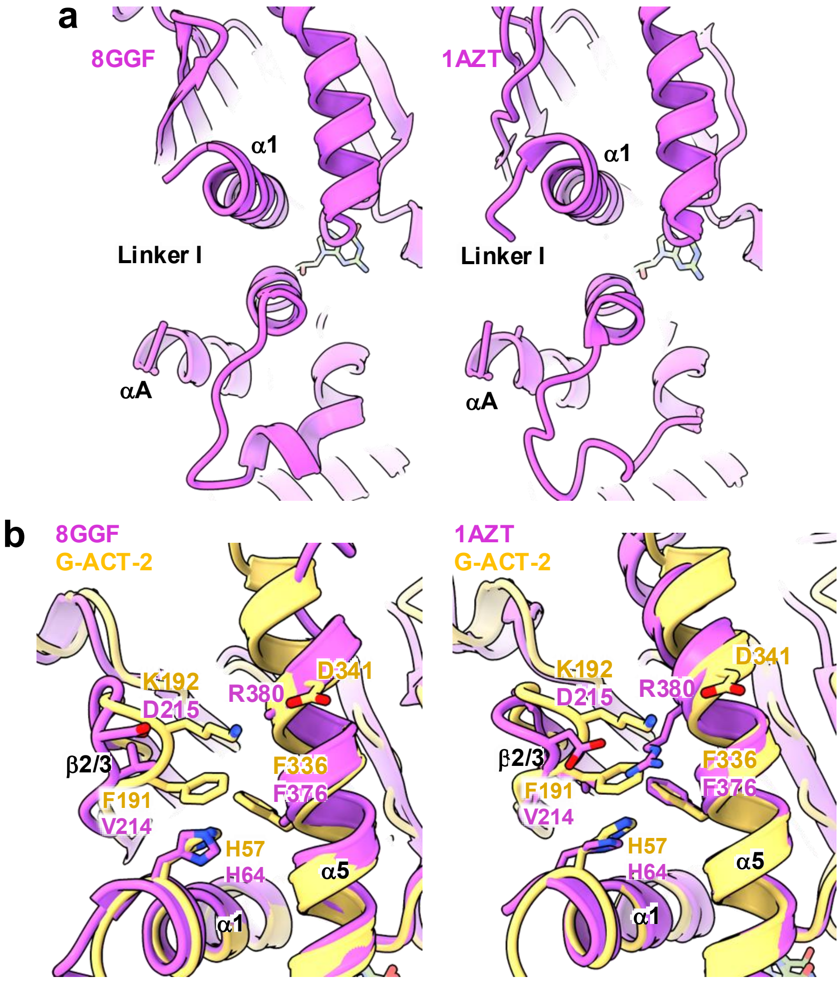


**Extended Data Figure 6: Structural differences between Gi and Gs a)** Comparison of the linker I region of Gs in the GTP-bound, cryo-EM frame 20 state (PDB:8GGF) and GTP-analogue bound crystal structure (PDB:1AZT). **b)** Comparison of Gi in the G-ACT-2 state with Gs in the GTP-bound, cryoEM frame 20 state (PDB:8GGF, left) and GTP-analogue bound crystal structure (PDB:1AZT, right).

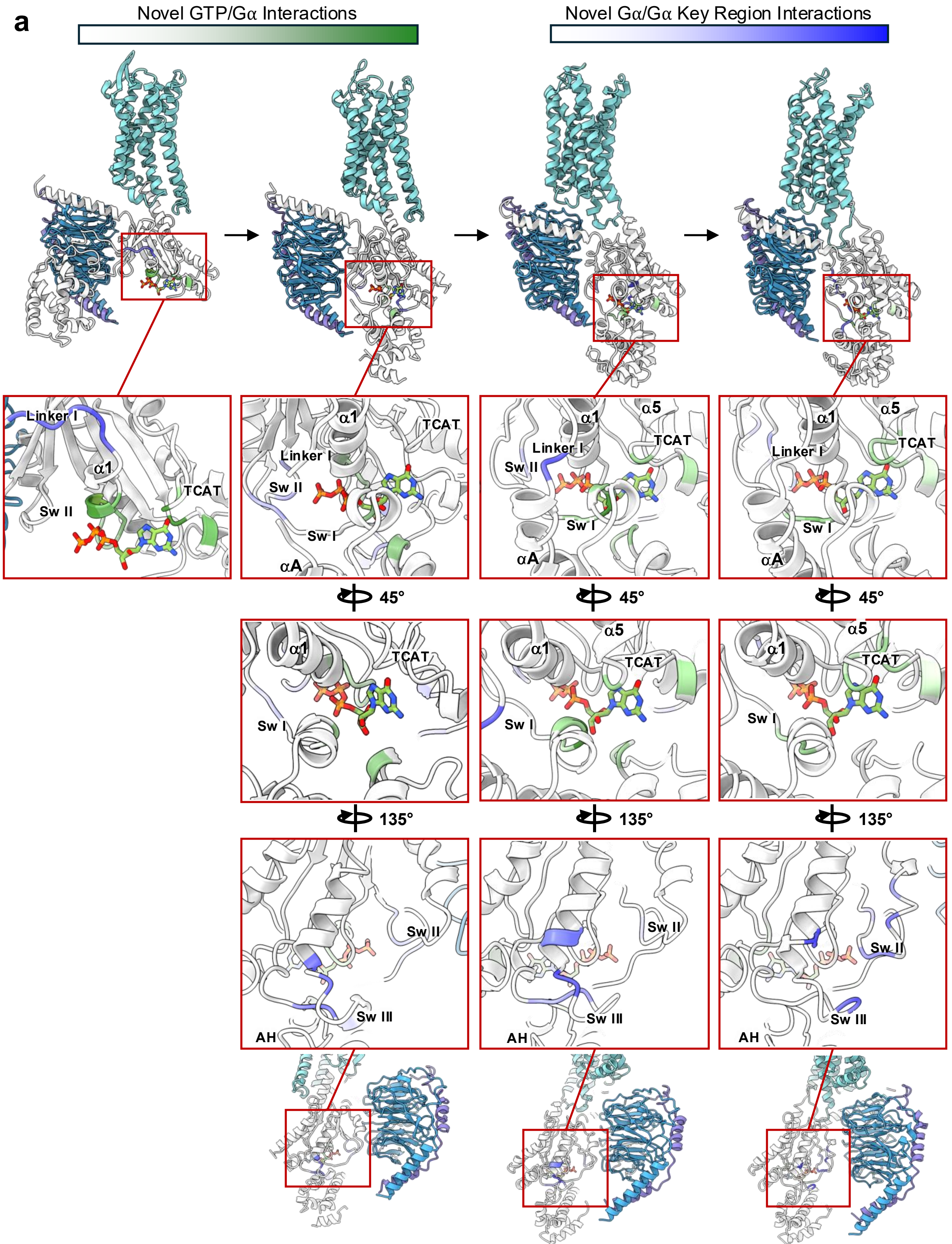


**Extended Data Figure 7: Novel interactions in the Gαi subunit during GTP-induced activation a)** Analysis of the formation of new GTP-Gα contacts (white-green scale) and new Gα-Gα contacts in the key linker I, switch I (linker II), switch II, and switch III regions (white-blue scale) as MOR/Gi GTP transitions from state G-Primed, G-ACT-1, G-ACT-2, and G-ACT-3.

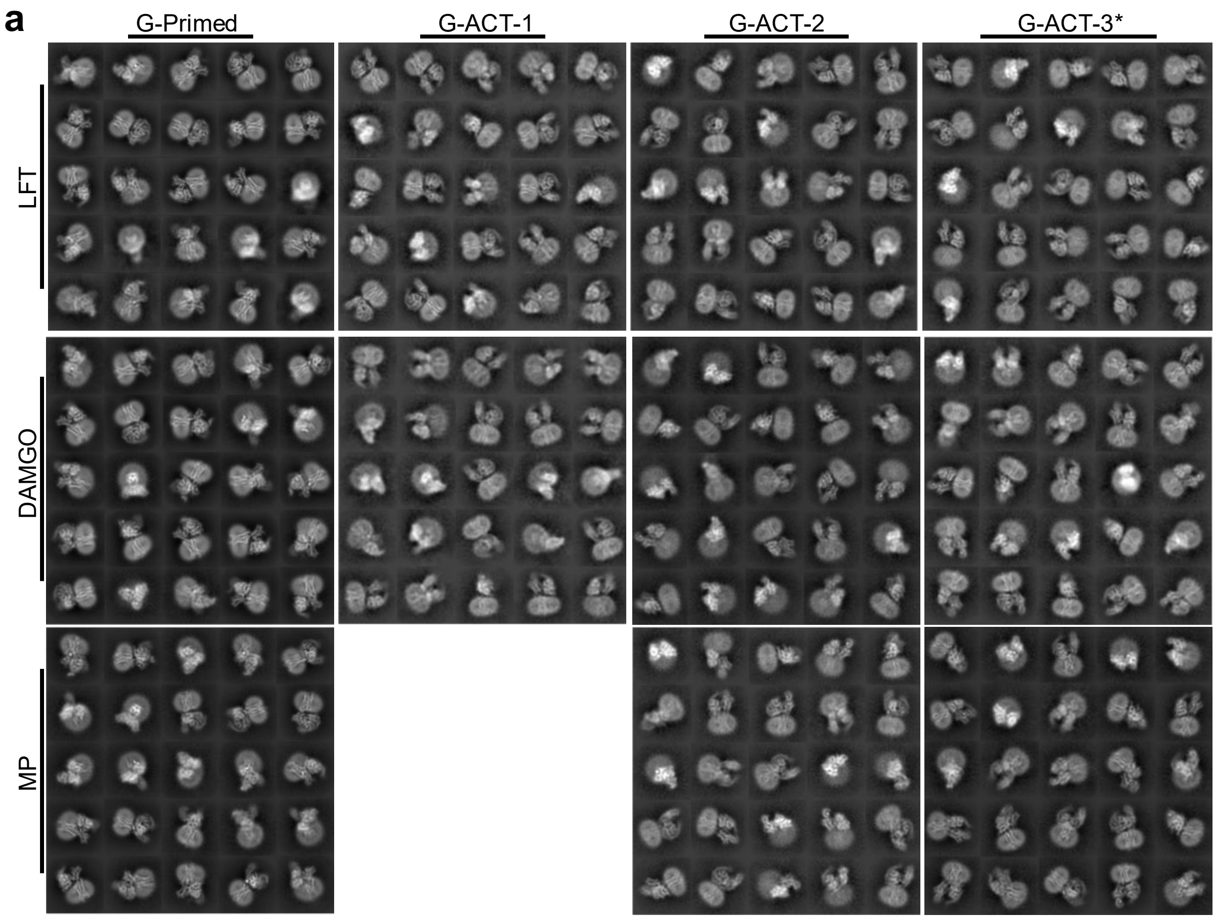


**Extended Data Figure 8: 2D Class averages for the main MOR/Gi baseline and intermediate states. a)** 2D Class Averages of all of the states resolved in this manuscript. G-ACT-3* corresponds to G-ACT-3 for MP and DAMGO, and G-ACT-2' for LFT.

| | No. Micrographs | Total Dose (e–/Å²) |
|---|---|---|
| **LFT** | | |
| Baseline | 4882 | 50 |
| 6s grid 1 | 7686 | 54 |
| 6s grid 2 | 14178 | 54 |
| 6s grid 3 | 6664 | 54 |
| 6s grid 4 | 9535 | 52 |
| 10s grid 1 | 6558 | 50 |
| 10s grid 2 | 9990 | 55 |
| 10s grid 3 | 12459 | 52 |
| 20s grid 1 | 10719 | 55 |
| 20s grid 2 | 9359 | 55 |
| 20s grid 3 | 7931 | 55 |
| 30s grid 1 | 13789 | 50 |
| 30s grid 2 | 12351 | 55 |
| 45s grid 1 | 15882 | 55 |
| **DAMGO** | | |
| Baseline | 6376 | 60 |
| 10s grid 1 | 4973 | 62 |
| 10s grid 2 | 5768 | 62 |
| 20s grid 1 | 11244 | 62 |
| 20s grid 2 | 7018 | 60 |
| 20s grid 3 | 6263 | 60 |
| 20s grid 4 | 9360 | 60 |
| 30s grid 1 | 13906 | 61 |
| 30s grid 2 | 3065 | 60 |
| 30s grid 3 | 8329 | 56 |
| 30s grid 4 | 2934 | 56 |
| 45s grid 1 | 8527 | 61 |
| 45s grid 2 | 7474 | 60 |
| 45s grid 3 | 7879 | 61 |
| **MP** | | |
| 10s grid 1 | 7974 | 55 |
| 10s grid 2 | 14206 | 55 |
| 10s grid 3 | 5602 | 55 |
| 30s grid 1 | 9745 | 55 |
| 30s grid 2 | 7428 | 55 |
| 30s grid 3 | 2840 | 55 |
| 45s grid 1 | 3208 | 55 |
| 45s grid 2 | 3506 | 55 |
| 45s grid 3 | 3777 | 55 |

**Extended Data Table 1:** Cryo-EM data sets collected in this work

| | MOR/Gi/ LFT Baseline Global (PDB:9ODE) (EMD-70356) | MOR/Gi/ LFT G-Primed Global (PDB:9ODF) (EMD-70357) | MOR/Gi/ LFT G-Primed Local MOR (EMD-70357) | MOR/Gi/ LFT G-ACT-1 Global (PDB:9ODG) (EMD-70358) | MOR/Gi/ LFT G-ACT-1 Gi Local (EMD-70358) | MOR/Gi/ LFT G-ACT-2/3 Global (EMD-70361) |
|---|---|---|---|---|---|---|
| **Data collection and processing** | | | | | | |
| Voltage (kV) | 300 | 300 | 300 | 300 | 300 | 300 |
| Electron exposure (e–/Å$^2$) | 50 | 50-55 | 50-55 | 50-55 | 50-55 | 50-55 |
| Defocus range (μm) | -0.6 to -1.8 | -0.6 to -1.8 | -0.6 to -1.8 | -0.6 to -1.8 | -0.6 to -1.8 | -0.6 to -1.8 |
| Pixel size (Å) | 0.96 | 0.96 | 0.96 | 0.96 | 0.96 | 0.96 |
| Symmetry imposed | C1 | C1 | C1 | C1 | C1 | C1 |
| Final particle images (no.) | 592,581 | 3,286,889 | 3,286,889 | 78,212 | 78,212 | 271,332 |
| Map resolution (Å) | 2.5 | 2.4 | 2.4 | 4.3 | 3.8 | 3.5 |
| FSC threshold | 0.143 | 0.143 | 0.143 | 0.143 | 0.143 | 0.143 |
| Map sharpening B factor (Å$^2$) | 80.2 | 97.7 | 97.7 | 171.4 | 126.8 | 168.2 |
| **Refinement** | | | | | | |
| Initial Model Used (PDB Code) | 7T2H | 7T2H | | 7T2H, 1GP2 | | |
| Model Resolution | 2.5 | 2.4 | | 4.2 | | |
| FSC Threshold | 0.143 | 0.143 | | 0.143 | | |
| *Model Composition* | | | | | | |
| Non-hydrogen | 6590 | 6614 | | 5842 | | |
| *B factor (Å$^2$)* | | | | | | |
| Protein Atoms | 50.6 | 34.7 | | 76.5 | | |
| Ligands | 73.4 | 48.8 | | 82.2 | | |
| *R.M.S Deviations* | | | | | | |
| Bonds (Å) | 0.006 | 0.004 | | 0.002 | | |
| Angles (º) | 0.950 | 0.751 | | 0.672 | | |
| *Validation* | | | | | | |
| MolProbity score | 1.50 | 1.80 | | 1.47 | | |
| Clashscore | 5.06 | 9.35 | | 4.24 | | |
| Poor rotamers (%) | 0.76 | 0.61 | | 0.00 | | |
| *Ramachandran Plot* | | | | | | |
| Favored (%) | 96.50 | 95.63 | | 96.10 | | |
| Allowed (%) | 3.39 | 4.26 | | 3.80 | | |
| Outliers (%) | 0.12 | 0.12 | | 0.10 | | |

| | MOR/Gi/ LFT G-ACT-2 G Local (PDB:9ODH) (EMD-70359) | MOR/Gi/ LFT G-ACT-2' G Local (PDB:9ODI) (EMD-70360) | MOR/Gi/ LFT G-ACT-2/3 G Local (EMD-70361) | MOR/Gi/ LFT G-ACT-2 Global (EMD-70362) | MOR/Gi/ LFT G-ACT-2' Global (EMD-70363) |
|---|---|---|---|---|---|
| **Data collection and processing** | | | | | |
| Voltage (kV) | 300 | 300 | 300 | 300 | 300 |
| Electron exposure (e–/Å2) | 50-55 | 50-55 | 50-55 | 50-55 | 50-55 |
| Defocus range (μm) | -0.6 to -1.8 | -0.6 to -1.8 | -0.6 to -1.8 | -0.6 to -1.8 | -0.6 to -1.8 |
| Pixel size (Å) | 0.96 | 0.96 | 0.96 | 0.96 | 0.96 |
| Symmetry imposed | C1 | C1 | C1 | C1 | C1 |
| Final particle images (no.) | 49,485 | 66,784 | 271,332 | 65,217 | 67,121 |
| Map resolution (Å) | 3.8 | 3.7 | 3.3 | 3.9 | 4.4 |
| FSC threshold | 0.143 | 0.143 | 0.143 | 0.143 | 0.143 |
| Map sharpening B factor (Å$^2$) | 112.9 | 107.6 | 135.3 | 135.9 | 164.7 |
| **Refinement** | | | | | |
| Initial Model Used (PDB Code) | 1GP2 | 1GP2 | | | |
| Model Resolution | 3.7 | 3.6 | | | |
| FSC Threshold | 0.143 | 0.143 | | | |
| *Model Composition* | | | | | |
| Non-hydrogen | 4395 | 4614 | | | |
| *B factor (Å$^2$)* | | | | | |
| Protein Atoms | 61.5 | 58.8 | | | |
| Ligands | 69.6 | 67.8 | | | |
| *R.M.S Deviations* | | | | | |
| Bonds (Å) | 0.003 | 0.004 | | | |
| Angles (º) | 0.753 | 0.838 | | | |
| *Validation* | | | | | |
| MolProbity score | 1.68 | 1.52 | | | |
| Clashscore | 5.97 | 5.18 | | | |
| Poor rotamers (%) | 1.18 | 0.00 | | | |
| *Ramachandran Plot* | | | | | |
| Favored (%) | 95.71 | 96.32 | | | |
| Allowed (%) | 4.14 | 3.53 | | | |
| Outliers (%) | 0.15 | 0.15 | | | |

**Extended Data Table 2:** Cryo-EM Map & Model Statistics, LFT Data

| | MOR/Gi/ MP G-Primed 3DVA Filtered (PDB:9ODJ) (EMD-70364) | MOR/Gi/ MP G-Primed Global (EMD-70367) | MOR/Gi/ MP G-ACT-2/3 Consensus (EMD-70368) | MOR/Gi/ MP G-ACT-2 Global (PDB:9ODK) (EMD-70365) | MOR/Gi/ MP G-ACT-2 Gi Local (EMD-70365) | MOR/Gi/ MP G-ACT-3 Global (PDB:9ODL) (EMD-70366) |
|---|---|---|---|---|---|---|
| **Data collection and processing** | | | | | | |
| Voltage (kV) | 300 | 300 | 300 | 300 | 300 | 300 |
| Electron exposure (e–/Å$^2$) | 55 | 55 | 55 | 55 | 55 | 55 |
| Defocus range (μm) | -0.6 to -1.8 | -0.6 to -1.8 | -0.6 to -1.8 | -0.6 to -1.8 | -0.6 to -1.8 | -0.6 to -1.8 |
| Pixel size (Å) | 0.96 | 0.96 | 0.96 | 0.96 | 0.96 | 0.96 |
| Symmetry imposed | C1 | C1 | C1 | C1 | C1 | C1 |
| Final particle images (no.) | 132,871 | 411,010 | 208,885 | 44,646 | 44,646 | 57,982 |
| Map resolution (Å) | 3.0 | 2.9 | 3.6 | 3.9 | 3.8 | 3.6 |
| FSC threshold | 0.143 | 0.143 | 0.143 | 0.143 | 0.143 | 0.143 |
| Map sharpening B factor (Å$^2$) | 83.6 | 96.9 | 147.0 | 116.2 | 93.6 | 120.9 |
| **Refinement** | | | | | | |
| Initial Model Used (PDB Code) | 7T2G | | | 1GP2 | | 7UL4, 1GP2 |
| Model Resolution | 3.0 | | | 3.8 | | 3.6 |
| FSC Threshold | 0.143 | | | 0.143 | | 0.143 |
| *Model Composition* | | | | | | |
| Non-hydrogen | 7558 | | | 6577 | | 7165 |
| *B factor (Å$^2$)* | | | | | | |
| Protein Atoms | 88.7 | | | 81.9 | | 107.7 |
| Ligands | 103.7 | | | 53.1 | | 105.5 |
| *R.M.S Deviations* | | | | | | |
| Bonds (Å) | 0.004 | | | 0.003 | | 0.003 |
| Angles (°) | 0.699 | | | 0.696 | | 0.631 |
| *Validation* | | | | | | |
| MolProbity score | 1.44 | | | 1.78 | | 1.53 |
| Clashscore | 5.13 | | | 8.23 | | 5.00 |
| Poor rotamers (%) | 0.97 | | | 0.21 | | 0.50 |
| *Ramachandran Plot* | | | | | | |
| Favored (%) | 97.03 | | | 95.14 | | 96.05 |
| Allowed (%) | 2.87 | | | 4.86 | | 3.95 |
| Outliers (%) | 0.10 | | | 0.00 | | 0.00 |

| | MOR/Gi/ MP G-ACT-3 Gi Local (EMD-70365) |
|---|---|
| **Data collection and processing** | |
| Voltage (kV) | 300 |
| Electron exposure (e–/Å2) | 55 |
| Defocus range (μm) | -0.6 to -1.8 |
| Pixel size (Å) | 0.96 |
| Symmetry imposed | C1 |
| Final particle images (no.) | 57,982 |
| Map resolution (Å) | 3.6 |
| FSC threshold | 0.143 |
| Map sharpening B factor (Å$^2$) | 88.4 |
| **Refinement** | |
| Initial Model Used (PDB Code) | |
| Model Resolution | |
| FSC Threshold | |
| *Model Composition* | |
| Non-hydrogen | |
| *B factor (Å$^2$)* | |
| Protein Atoms | |
| Ligands | |
| *R.M.S Deviations* | |
| Bonds (Å) | |
| Angles (°) | |
| *Validation* | |
| MolProbity score | |
| Clashscore | |
| Poor rotamers (%) | |
| *Ramachandran Plot* | |
| Favored (%) | |
| Allowed (%) | |
| Outliers (%) | |

**Extended Data Table 3:** Cryo-EM Map & Model Statistics, MP Data

| | MOR/Gi/ DAMGO Baseline Global (PDB:9ODM) (EMD-70369) | MOR/Gi/ DAMGO Baseline MOR Local (EMD-70369) | MOR/Gi/ DAMGO G-Primed Global (PDB:9ODN) (EMD-70370) | MOR/Gi/ DAMGO G-Primed AHD Filtered (EMD-70370) | MOR/Gi/ DAMGO G-ACT-1 Global (EMD-70373) | MOR/Gi/ DAMGO G-ACT-2/3 Consensus (EMD-70374) |
|---|---|---|---|---|---|---|
| **Data collection and processing** | | | | | | |
| Voltage (kV) | 300 | 300 | 300 | 300 | 300 | 300 |
| Electron exposure (e–/Å$^2$) | 60 | 60 | 56-62 | 56-62 | 56-62 | 56-62 |
| Defocus range (μm) | -0.6 to -1.8 | -0.6 to -1.8 | -0.6 to -1.8 | -0.6 to -1.8 | -0.6 to -1.8 | -0.6 to -1.8 |
| Pixel size (Å) | 0.8677 | 0.8677 | 0.8677 | 0.8677 | 0.8677 | 0.8677 |
| Symmetry imposed | C1 | C1 | C1 | C1 | C1 | C1 |
| Final particle images (no.) | 394,504 | 394,504 | 2,756,068 | 854,816 | 56,870 | 198,050 |
| Map resolution (Å) | 2.8 | 3.1 | 2.4 | 2.7 | 6.5 | 3.9 |
| FSC threshold | 0.143 | 0.143 | 0.143 | 0.143 | 0.143 | 0.143 |
| Map sharpening B factor (Å$^2$) | 100.2 | 92.0 | 94.1 | 93.5 | 403.4 | 183.2 |
| **Refinement** | | | | | | |
| Initial Model Used (PDB Code) | 6DDE | | 6DDE | | | |
| Model Resolution | 2.8 | | 2.4 | | | |
| FSC Threshold | 0.143 | | 0.143 | | | |
| *Model Composition* | | | | | | |
| Non-hydrogen | 6483 | | 6479 | | | |
| *B factor (Å$^2$)* | | | | | | |
| Protein Atoms | 38.4 | | 33.8 | | | |
| Ligands | 54.6 | | 46.6 | | | |
| *R.M.S Deviations* | | | | | | |
| Bonds (Å) | 0.004 | | 0.007 | | | |
| Angles (°) | 0.682 | | 0.982 | | | |
| *Validation* | | | | | | |
| MolProbity score | 1.31 | | 1.39 | | | |
| Clashscore | 3.99 | | 2.89 | | | |
| Poor rotamers (%) | 0.81 | | 0.82 | | | |
| *Ramachandran Plot* | | | | | | |
| Favored (%) | 97.34 | | 95.53 | | | |
| Allowed (%) | 2.66 | | 4.47 | | | |
| Outliers (%) | 0.00 | | 0.00 | | | |

| | MOR/Gi/ DAMGO G-ACT-2 Global (PDB:9ODO) (EMD-70371) | MOR/Gi/ DAMGO G-ACT-2 G Local (EMD-70371) | MOR/Gi/ DAMGO G-ACT-3 Global (PDB:9ODP) (EMD-70372) | MOR/Gi/ DAMGO G-ACT-3 G Local (EMD-70372) |
|---|---|---|---|---|
| **Data collection and processing** | | | | |
| Voltage (kV) | 300 | 300 | 300 | 300 |
| Electron exposure (e–/Å2) | 56-62 | 56-62 | 56-62 | 56-62 |
| Defocus range (μm) | -0.6 to -1.8 | -0.6 to -1.8 | -0.6 to -1.8 | -0.6 to -1.8 |
| Pixel size (Å) | 0.8677 | 0.8677 | 0.8677 | 0.8677 |
| Symmetry imposed | C1 | C1 | C1 | C1 |
| Final particle images (no.) | 52,140 | 52,140 | 45,305 | 45,305 |
| Map resolution (Å) | 3.8 | 3.6 | 4.3 | 4.2 |
| FSC threshold | 0.143 | 0.143 | 0.143 | 0.143 |
| Map sharpening B factor (Å$^2$) | 135.0 | 129.6 | 150.6 | 148.3 |
| **Refinement** | | | | |
| Initial Model Used (PDB Code) | 1GP2 | | 1GP2 | |
| Model Resolution | 3.6 | | 4.1 | |
| FSC Threshold | 0.143 | | 0.143 | |
| *Model Composition* | | | | |
| Non-hydrogen | 4586 | | 4248 | |
| *B factor (Å$^2$)* | | | | |
| Protein Atoms | 91.1 | | 113.9 | |
| Ligands | 93.9 | | 103.8 | |
| *R.M.S Deviations* | | | | |
| Bonds (Å) | 0.003 | | 0.002 | |
| Angles (°) | 0.556 | | 0.532 | |
| *Validation* | | | | |
| MolProbity score | 1.61 | | 1.18 | |
| Clashscore | 5.12 | | 2.89 | |
| Poor rotamers (%) | 0.81 | | 0.00 | |
| *Ramachandran Plot* | | | | |
| Favored (%) | 95.12 | | 97.48 | |
| Allowed (%) | 4.88 | | 2.52 | |
| Outliers (%) | 0.00 | | 0.00 | |

**Extended Data Table 4:** Cryo-EM Map & Model Statistics, DAMGO Data
